# Modelling the Replay of Dynamic Memories from Cortical Alpha Oscillations with the Sync-Fire / deSync Model

**DOI:** 10.1101/2020.01.28.921205

**Authors:** George Parish, Sebastian Michelmann, Simon Hanslmayr, Howard Bowman

**Affiliations:** School of Psychology and Centre for Human Brain Health, University of Birmingham, UK.; Department of Computer Science, University of Kent, UK.; Department of Psychology, Princeton University, US.

## Abstract

We here propose a neural network model to explore how neural oscillations might regulate the replay of memory traces. We simulate the encoding and retrieval of a series of events, where temporal sequences are uniquely identifiable by analysing population activity, as several recent EEG/MEG studies have shown. Our model comprises three parts, each considering distinct hypotheses. A cortical region actively represents sequences through the disruption of an intrinsically generated alpha rhythm, where a desynchronisation marks information-rich operations as the literature predicts. A binding region converts each event into a discrete index, enabling repetitions through a sparse encoding of events. We also instantiate a temporal region, where an oscillatory “ticking-clock” made up of hierarchical synfire chains discretely indexes a moment in time. By encoding the absolute timing between events, we show how one can use cortical desynchronisations to dynamically detect unique temporal signatures as they are replayed in the brain.

## INTRODUCTION

There has been much work in recent years to understand how the brain represents ‘what’ and ‘when.’ With respect to the ‘what’ of episodic memory, neural oscillations are thought to play a key role in structuring our informational capacity (Fell & Axmacher, 2011). It has been proposed that information flow depends on whether neural activation is restricted to the facilitatory phase of an entraining oscillation or not, such that cortical brain oscillations would thus provide a gating function to the representation of information (Klimesch, et al., 2007; Hanslmayr, et al., 2012; Parish, et al., 2018; Jensen & Mazaheri, 2010). If this were the case, information representation would be measurable by oscillatory power changes in specific frequencies. As such, alpha frequency de-synchronisations are thought to signify information processing in cortical regions, as de-regulated neural activation enables potential information gain (Hanslmayr, et al., 2012). It has also been shown that successful episodic memory encoding and retrieval can be predicted by alpha power in several experimental (Fell, et al., 2011; Hanslmayr & Staudigl, 2014; Hanslmayr, et al., 2012; Khader, et al., 2010; Klimesch, et al., 2005; Waldhauser, et al., 2016), and modelling studies (Parish, et al., 2018).

Where previous computational work has shown that alpha power decreases can signal high information states (Parish, et al., 2018), the current modelling work aims to relate the phase of on-going alpha to information content. We here employ a method, representational similarity analysis (RSA), which has recently been employed to detect the active representation of neural patterns at encoding (Ng, et al., 2013; Schyns, et al., 2011) and retrieval (Staudigl, et al., 2015; Staresina, et al., 2016; Wimber, et al., 2012; Yaffe, et al., 2014), based upon phase-reset oscillatory signatures. More recent studies have even associated these temporal signatures with specific stimuli, enabling RSA to dynamically decode which temporal pattern is being actively replayed by the brain, and when (Michelmann, et al., 2016; Michelmann, et al., 2018; Michelmann, et al., 2019). This RSA method has been applied to detect replay of visual or auditory stimuli (Staudigl & Hanslmayr, 2019), replay of complex movie sequences (Michelmann, et al., 2019), replay in working memory and episodic memory (Michelmann, et al., 2018), and even replay during sleep (Schreiner, et al., 2018). On a more general level, a number of recent MEG studies leveraged other multivariate analysis methods to detect replay of sequences (Kornysheva, et al., 2019; Kurth-Nelson, et al., 2016; Lui, et al., 2019), which raises the question of what the neural mechanisms are that give rise to the detection of sequential replay in a large scale population signal as recorded with EEG/MEG. We therefore intend to test our theorised temporal replay framework by reproducing the findings of a recent episodic memory paradigm defined by Michelmann, et al., 2016, where unique dynamic stimuli (audio & video) are successfully identified at retrieval using their sequential oscillatory signatures at encoding.

With respect to the ‘when’ of episodic memory, paradigms have demonstrated that humans can integrate experiences in time and space within the realm of milliseconds to minutes (Mauk & Buonomano, 2004; Tsao, et al., 2018). This is thought to be mediated by the hippocampus within the medial-temporal-lobe (MTL) (Howard & Eichenbaum, 2015; O’Reilly, et al., 2011; Squire, 1992; Squire, et al., 2004). Here, a gradually changing state of temporal context is thought to serve as a cue for episodic recall (Howard & Eichenbaum, 2013; Howard, et al., 2012), as theorised by other models of working memory (Burgess & Hitch, 1999; Shankar & Howard, 2012). It has also been hypothesised that there is a competitive architecture underpinning spatial working memory, navigation and path integration in the hippocampus (see McNaughton, et al., 2006, for review). The same might be said for the timing aspect of episodic memory, where studies have indicated that the hippocampus holds many temporal sequences that it can successfully associate, disambiguate and navigate between (MacDonald, et al., 2013; Pastalkova, et al., 2008; Wood, et al., 2000). Competition is often modelled as short-range excitatory and long-range inhibitory connections in what is often termed a ‘Mexican-hat’ topology. Together, this encourages winner-take-all behaviour, an architecture that has commonly been used in hierarchical models of vision, recognition and attention (Carpenter & Grossberg, 1987; Itti, et al., 1998; Reisenhuber & Poggio, 1999). A hypothesis of this modelling work is that any model of temporal sequencing would enable competition through a winner-take-all architecture, similar to other temporal sequencing models (Itskov, et al., 2011; Rolls & Mills, 2019), whereby the emergence of a winning pathway maintains temporal context through the suppression of losing pathways.

The encoding of temporal sequences is thought to be enabled by the sequential activation of time cells within the hippocampus (see Eichenbaum, 2014, for review), which have been theorised to exist as a series of feed-forward chains, typically called synfire chains, in a previous modelling study (Itskov, et al., 2011). Feed-forward activation is thought to be the best enabler for fast communication with high temporal precision, thought necessary for the distributed time-keeping that occurs during sensory and motor events (Mauk & Buonomano, 2004), possibly by enabling target cells to conjunctively represent distinct elements of a stimulus (Fries, 2005). Models of such synfire chains have shown that this kind of signal transmission can operate within a noisy environment (Diesmann, et al., 1999), can co-exist within a randomly connected embedding network (Kumar, et al., 2008), and might even naturally emerge in a plastic and locally connected environment (Fiete, et al., 2010). It can be further demonstrated that a feed-forward architecture can support temporal encoding (Itskov, et al., 2011), predicting that temporal sequences can be internally generated, being reliable from trial to trial, context dependent and long lasting, in a manner similar to time cells (Eichenbaum, 2014). However, a common criticism of such a means to model time is that the length of the chain must increase linearly with the desired duration (Shankar & Howard, 2012). This complexity issue might be further compounded by the aforementioned observations that there might be many competing time-cell chains within the hippocampus (MacDonald, et al., 2013; Pastalkova, et al., 2008; Wood, et al., 2000).

A fundamental goal of the temporal aspect of the modelling reported here is to thus reduce the physical requirements of synfire chains in the encoding of longer temporal durations. We do this by instantiating a hierarchical feed-forward chain, echoing principles of a recently published model (Rolls & Mills, 2019), where sequential cell assemblies maintain persistent activation in a feed-forward manner (Goldman, 2009). To do this, we envisage a simple cell assembly of Hodgkin-Huxley neurons that can be scaled up to encode for multiple, simultaneous and interacting temporal hierarchies. This cellular assembly exists as a compact unit with specific feedforward and feedback connections, initiating and terminating persistent activation upon the completion of hierarchical sequences. We thus instantiate a hierarchy of synfire chains, where completion of a lower-order, faster temporal dimension initiates the transmission of persistent activity to the next node in a higher-order, slower temporal dimension. As such, lower-order sequences repeat at every node of the higher-order sequence, decreasing the physical requirements of feedforward chains in the encoding of long temporal durations. A moment in time is then marked as the concurrent activation of multiple temporal positions on simultaneous hierarchies, much like the multi-dimensional hands of a ticking clock, as shown in the video of Figure 1. As a recent model has theorised (Rolls & Mills, 2019), hierarchies of networks with varying time constants might be linked to time ramping cells in the lateral entorhinal cortex (Kropff & Treves, 2008), where synchrony across networks enables robust and reliable temporal coding.

**FIGURE 1.** SEE VIDEO FILE SYNFIRE CHAINS.MP4. Hierarchical updates between time-keepers of varying temporal dimensions. Excitatory (lines with arrow-heads) and inhibitory (lines with circle-heads) connections dictate how neural assemblies (grey circles) generally interact (see supplementary Figure S4 for a more precise description of feed-forward and feed-back connectivity). Highlighted circles and lines indicate currently active assemblies and pathways, respectively. Several temporal layers simultaneously encode for the present moment. Transiently active groups pass activation forwards (clockwise) through a synfire chain. Upon completion of any lower-order chain (1^st^ layer, inner circle), a feedback signal is sent to any higher-order layers (2^nd^ layer, outer circle) with the effect of terminating persistent activation in the active group of that layer. This process is shown in the descriptive video, where feedforward initiation from higher-to-lower order layers and feedback termination from lower-to-higher order layers organises feed-forward signal transmission.

Interestingly, in the application of a hierarchical encoding of time, it was found that events were bound with fewer errors if temporal synchronisation was somewhat oscillatory (see Figure 2 for description of “temporal conjunction errors”). This suggests the role that oscillations might play in segregating temporal episodes, as previous models have suggested (Brown, et al., 2000; Jensen, et al., 1996). As such, we liken our hierarchical synfire chain to the hierarchy of nested oscillatory frequencies that are thought to control neuronal excitability and stimulus processing in the auditory cortex (Lakatos, et al., 2005), and which are also present in the hippocampus during all manner of learning paradigms (Buzsáki, 2002; Fell & Axmacher, 2011). In this sense, we here initiate an oscillatory clock-time that is capable of defining precise inter-stimulus temporal periods in the manner of a hierarchically ticking clock. Our generation of sequence is thus similar to that of Jensen, et al. (1996), where nested oscillatory theta and gamma frequencies were theorised to maintain ordinal sequences, though here we theorise that they might also play a role in maintaining an absolute clock-like rhythm.

**FIGURE 2.**
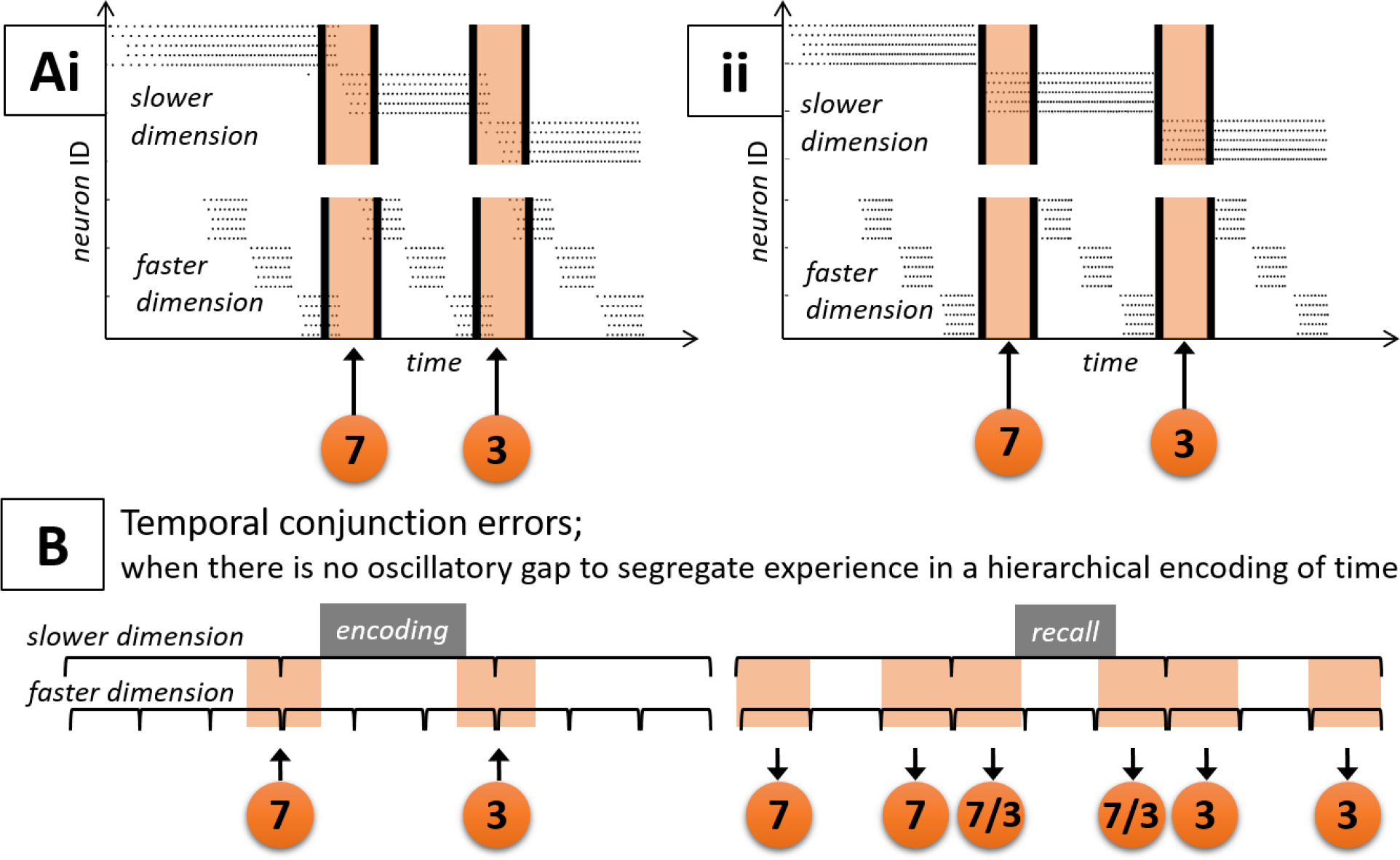
**A** The interaction between a feedback-terminating higher order chain (slower dimension) and a self-completing lower-order chain (faster dimension) is shown in this raster plot, where each spike event is represented by a black dot. This example simulation chose 3 neuronal groups per dimension, though this number can be scaled up – as later simulations will demonstrate. Here, two separate parameter sets are shown **(Ai-ii)**, which led to separate scenarios. In the first **(Ai)**, feed-forward transmission between hierarchies is immediate, meaning that items (orange circles) may be bound to multiple groups within each temporal dimension (indicated by orange shaded regions). This in turn might have the effect of causing order-errors or conjunction errors in the encoding of episodic sequences (Botella, et al., 1992). For example, in panel **B**, the hypothetical items 7 & 3 are bound to multiple groups in both temporal dimensions (**B**, left; indicated by orange shaded regions that overlap multiple slots), meaning that they are incorrectly recalled multiple times (**B**, right; indicated by orange shaded regions in multiple slots). This effect, that we call a “temporal conjunction error,” might arise if a clock-like mechanism is involved in the encoding of sequences. However, we show how one can minimise these errors by altering the parameter set slightly. In the simulation of panel **Aii**, one is left with a ‘hand-over’ period (indicated by orange shaded regions), where ramping activation is required prior to feed-forward transmission. During this time, incoming items (orange circles) might not be fully encoded, though the risk of temporal conjunction errors is minimalised. One could assume this issue to be significant for any hierarchical method that might be used to maintain temporal rhythm, suggesting a functional role for nested oscillations in the segregation of temporal episodes.

## MODEL ARCHITECTURE

Our model comprises three distinct mechanisms, as shown in Figure 3, where we aim to provide evidence for the inhibition-timing and information via desynchronisation hypotheses (Klimesch, et al., 2007; Hanslmayr, et al., 2012) by reproducing an experiment that decodes temporal signatures from neocortical alpha phase patterns (Michelmann, et al., 2016). As such, information representation occurs in a neocortical (NC) area (Figure 3, top section), where an alpha desynchronisation indicates information processing (Hanslmayr, et al., 2012), as described in previous modelling work (Parish, et al., 2018). In the current work, we create an intrinsic and dynamic alpha frequency through recurrent excitatory-inhibitory interactions (Figure 3A, top section; blue excitatory node & red inhibitory node). This allows changes in the environment to cause phase reset patterns with some specific temporal pattern (Figure 3B, top section; phase angle time series punctuated by event-driven phase-resets), thus conveying temporal information (Michelmann, et al., 2016; Ng, et al., 2013; Schyns, et al., 2011).

**FIGURE 3.**
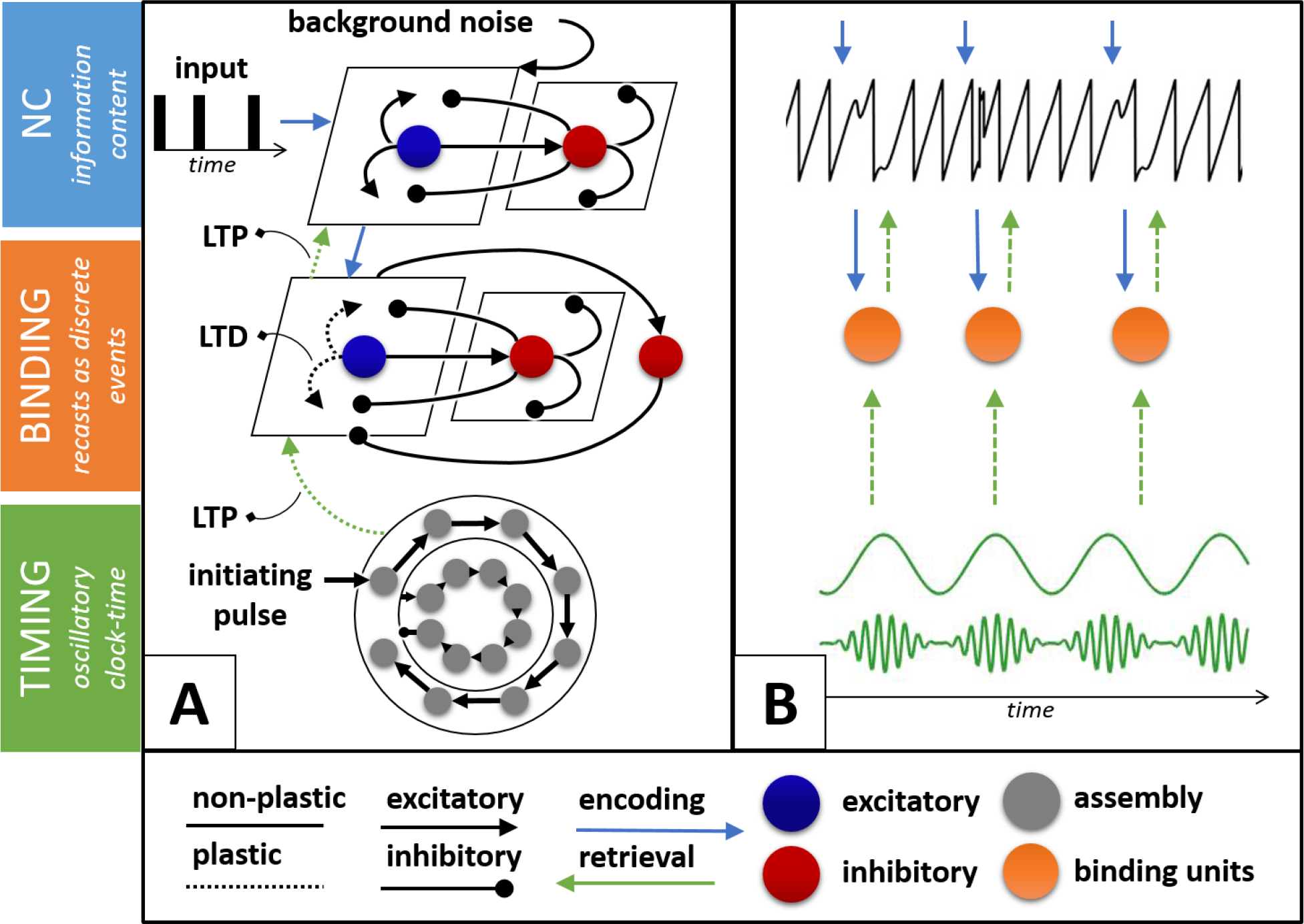
Architecture of a spiking neural network model, using Hodgkin-Huxley equations (Hodgkin & Huxley, 1952). A neo-cortical (NC) region made up of recurrent excitatory-inhibitory interactions, is made to intrinsically oscillate through the reception of low-level (non-oscillatory) background noise (**A**, top section: blue excitatory population & red inhibitory population). The selectiveness of this region, i.e. the “winner-take-all” behaviour encouraged by short-range excitatory and long-range inhibitory connections, enables the clear representation of each input, which in turn ensures a population phase-reset. During encoding (**A**; cyan top-down arrows), a sequence of incoming cortical stimuli trigger activation in a binding pool (**A**, middle section: blue excitatory population, red inhibitory population & additional red inhibitory “off-switch” node), a highly selective population that indexes each stimulus as a unique event through a combination of long-term potentiation (LTP) and hetero-synaptic long-term depression (LTD). Due to a highly selective parameter space, an additional inhibitory node was required to prevent runaway excitation, what we term an “off-switch”. Through LTP, active binding assemblies form a direct connection between concurrently active hierarchical synfire chains (**A**, bottom section: a feed-forward & clock-like structure used for the encoding of time, described in Figure 1), and the NC representation of the stimulus. Conversely, LTD occurs within active binding pool assemblies, diminishing the likelihood that they would be able to compete for the indexing of subsequent stimuli. To simplify our model, we here adapt our STDP rule by shifting the parameter subspace such that only LTD can occur between active binding units, though in reality we imagine this process to be caused by hetero-synaptic LTD in response to the LTP occurring on other dendrites (Volgushev, et al., 2016). Altogether, binding pool LTP & LTD ensures a sparse coding to index unique events. During recall (**A**, green bottom-up arrows), synfire chains are re-started and the relevant bindings are activated in sequence until the original pattern is re-instantiated in representational regions. Visualising this process through time in panel **B**, observable phase-reset patterns emerge in the intrinsically oscillating cortical region to represent information content (**B**, top section: cyan top-down arrows indicate the occurrence of stimuli and subsequent phase-resets). Describing the top-down encoding state, a sparse coding then indexes events in the binding pool region (**B**, middle section: orange nodes indicate the occurrence of a unique binding assembly), which are bound to the ticking hierarchical oscillator (**B**, bottom section: nested oscillatory frequencies). This enables events to be bound relative to a temporal rhythm (**B**; cyan top-down arrows), ensuring that they can be recalled (**B**; green bottom-up arrows) with the correct absolute timing between events. We assume the presence of a neuro-modulator that switches information processing between the encoding direction (cyan top-down arrows) and the retrieval direction (green bottom-up arrows), which is important to prevent cross-communication from contaminating encoding and recall processes. This is achieved by setting the weights of each pathway to zero at the appropriate processing phase. See supplementary section 1 for more information on topology and parameter definitions.

During encoding (Figure 3A; cyan top-down arrows), event-triggered activation is forwarded to a binding pool area. A binding pool has been thought to be necessary to allow for repetition, where any repeated stimulus can be re-represented by a unique tag (Bowman & Wyble, 2007), allowing us to differentiate between repeating stimuli in a sequence. The occurrence of a stimulus is treated as a distinct event through the activation of a selective population of units in a winner-take-all environment (Figure 3A, middle section), implemented as long-range inhibitory and short-range excitatory connections (commonly termed a “Mexican-hat” topology”). In order to prevent runaway excitation in this highly selective population, an additional inhibitory group was required. This received slow ramping excitatory input, eventually clamping down on the whole excitatory population. As such, we define this additional inhibitory group as “off-switch” cells. During the binding process, a group of unique binding pool units are associated to any active cortical unit via a calcium dependent learning rule (Graupner & Brunel, 2012). As their local weights decrease during this process, which effectively models a form of hetero-synaptic long-term-depression (Volgushev, et al., 2016), active binding groups diminish the likelihood that they will be able to compete during successive events. Thus, active bindings are unique to the bound event. Concurrently active temporal units (Figure 3A, bottom section) are also associated with binding pool units, thus sequencing the timing of events as they occur. Once the hierarchical time-keepers are re-started in a cued-recall paradigm, the relevant binding pool units become active at specific moments in time (Figure 3A; green bottom-up arrows), causing a temporal pattern of events to be re-instantiated in the neo-cortex.

The model was used to simulate an experimental paradigm that could detect the content-specific reinstatement of temporal patterns in the neocortex (see Michelmann, et al., 2016, for experiment). In the experiment, several dynamic stimuli were presented to the participant (i.e. a video or sound), which were each associated to a non-dynamic stimulus (i.e. a word). Alpha frequencies were interrogated during the presentation of each dynamic stimulus at encoding; a period of post-stimulus alpha desynchronisation was observed that is typical of active neuronal assemblies (Haegens, et al., 2011; Hanslmayr, et al., 2011a). During this period, the phase angle pattern was observed, which has been shown to reliably convey information content (Ng, et al., 2013; Schyns, et al., 2011), where a window indicating a period of phase-resets was chosen for each pattern (similar to Figure 3B, top section). Later, the associated word was presented to the participants, where they were asked to remember the associated dynamic stimulus. Similarly, the phase angle pattern during a period of alpha de-synchronisation was observed. As the experimenters had no knowledge of the specific time that the participant internally recollected the dynamic stimulus, they had to dynamically find the period of time that the video was being mentally re-played. This was done using a convolution approach (see Figure 4C for method), where the consistency of phase angle difference over time was calculated, between phase-reset patterns at encoding and a window of equal length slid over a phase-angle time series at recall. This approach gave a dynamic similarity signal through time, where an increase signalled high similarity between the phase-reset pattern at encoding and retrieval, thus indicating the re-instatement of neural patterns. Further, one could find the difference between similarity measures of trials of the same stimulus and trials of different stimuli to determine a content specificity measure. This was found to be positive, indicating that dynamic stimuli could be identified by a unique temporal signature. We aim to replicate these findings by encoding and retrieving multiple trials of several “bar-code” like sequences to our model, before employing representational similarity analysis to determine whether each pattern could be successfully identified by a unique temporal signature from simulated cortical representations.

**FIGURE 4.**
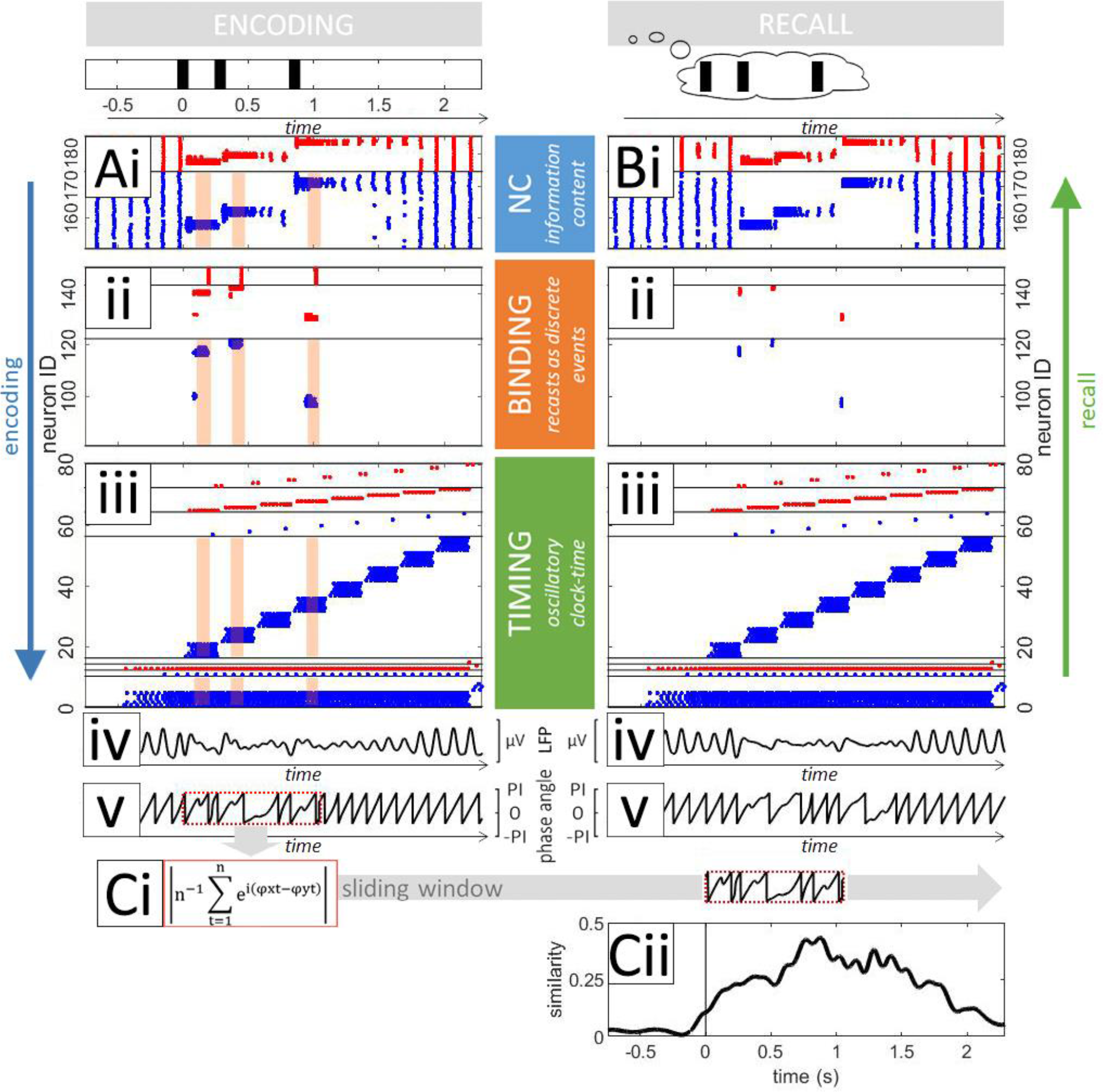
Raster plot of a single simulation (**A-B, i-iii**; blue/red dots = excitatory/inhibitory spike events) and subsequent phase-angle RSA (**iv**; local field potential (LFP), **v**; phase-angles, **Ci**; RSA method and application, **Cii**; similarity signal through time), for encoding (**A**; left panels, see supplementary Figure S5 for more detailed information on binding processes at encoding) and recall (**B**; right panels, see supplementary Figure S6 for more detailed information on intrinsic properties of the network at recall). At encoding, a sequence of events (top left panel; black lines) were fed into a random selection of cortical excitatory units (**Ai**), which became active at the expense of the population due to a “Mexican-hat” topology. This triggered activation in a unique group of selective binding pool units (**Aii**), which was itself terminated by an additional “off-switch” node to prevent run-away excitation (red spike events, top line-separated panel). During this time, hierarchical synfire chains maintained temporal rhythm through sequential activation of cellular assemblies (**Aiii**). Long-term-potentiation (LTP) worked to bind any active units together in time, from the temporal region to the binding pool region, and from the binding pool region to the cortical region. Long-term depression (LTD) worked to reduce the weights of active binding assemblies, ensuring a sparse coding for each incoming cortical stimulus. Orange shaded regions are applied to the raster plots at encoding (**Ai-iii**) to visualise periods of synaptic modification, where calcium amplitudes were above the threshold for potentiation. During recall (**B**; right panels), the re-started temporal region replays relevant bound events in the correct temporal order. The disruption of the intrinsic cortical oscillation can be seen in the LFP (**A-B iv**), where a diminution signals a desynchronisation in the dominant frequency (set to ∼8Hz in this simulation). A subsequent phase-reset pattern at encoding is highlighted in the red dotted box (**Ci**), which was used in an RSA approach to detect the re-instatement of a similar phase-reset pattern at retrieval (**Cii**).

## RESULTS

A single trial of our simulated paradigm can be seen in Figure 4. Here, neo-cortical (NC) assemblies were designed such that they intrinsically oscillated at a particular frequency, determined by parameters for excitation/inhibition interactions (Brunel, 2000). In addition, a winner-take-all environment was instantiated, as has been theorised in models of visual working memory (Itti, et al., 1998). Hypothetically, this ensures that only one locally connected neuronal group can be active at any one time, minimising the simultaneous activation of multiple representations in a distributed manner. As can be seen in Figure 4Ai, parameters were chosen such that activation could spread linearly across the entire excitatory population before inhibitory interactions had time to clamp down, thus allowing intrinsic oscillations to emerge. Once input was fed into a group of excitatory units (Figure 4Ai; sustained excitatory spike events, depicted as blue dots), the selective topology can be seen inhibiting competing representations (Figure 4Ai; sustained inhibitory spike events, depicted as red dots). This subsequently causes a desynchronisation in the ongoing alpha oscillation (Figure 4Aiv; local field potential), a phenomenon shown in many studies (Haegens, et al., 2011; Hanslmayr, et al., 2011a; Klimesch, et al., 2007) and models (Parish, et al., 2018) to be related to neuronal activation. During event-related desynchronisations in cortical alpha oscillations, evaluating the phase-angle time-series has been shown to render qualitative information about stimulus content (Ng, et al., 2013; Schyns, et al., 2011). Therefore, in the model, we show that the phase-angle time-series does indeed show evidence of phase-resets at this time (Figure 4Av; red-dotted box), as the on-going intrinsic oscillation is externally affected through its representation of incoming stimuli. This indicates that these phase-reset periods are linked to periods of event-related cortical activation, lending theoretical evidence to the argument that phase-angle time-series can be used to decode information content (Ng, et al., 2013; Schyns, et al., 2011).

The binding pool within the model was developed with similar intentions as other models (Bowman & Wyble, 2007), where a unique node would activate to encode an incoming discrete event. It is argued that this method allows for repetitions, as each presentation is treated as an independent event. In this way, a binding node is only required to activate during an event-related cortical activation, as can be seen in the raster plot of Figure 4Aii. Here, parameters were chosen such that intrinsic oscillatory activity is not sufficient to cause activation in the binding pool. This is fulfilled by a relatively large synaptic time constant to gate cortical-binding projections, requiring sustained input to trigger binding activity. It was also important to obtain weight variation on these projections by sampling from a normal distribution, to increase the likelihood of winner-take-all behaviour during event-related sustained input. As depicted in Figure 3A, a global “off-switch” inhibitory cell adds an additional safeguard mechanism to prevent runaway activation across the excitatory population, operating to inhibit the entire binding pool once sustained excitatory input reaches a threshold. Active binding pool units engage in learning, making bindings between the active cortical content layer and the currently active temporal units that together, index a moment in time (indicated by orange shaded regions in Figure 4Ai-iii, see also supplementary Figure S5 for full analysis of weight change in the model). During this time, the connections between active binding pool assemblies undergo long-term-depression, meaning that these bound units would not be able to compete upon successive activations of the binding pool due to its highly selective topology. This ensured that each binding pool assembly uniquely indexed an event in time.

Figure 4Aiii also shows a raster plot of our hierarchical, clock-like synfire chains, similar to Figure 2A though here with more sequential groups. Once these chains are re-started with an initiating burst in Figure 4Biii, the relevant binding and cortical clusters are then successively activated dependent on the combined activation of synfire hierarchies (Figure 4Bi & 4Bii; see also supplementary Figure S6 for full analysis of synaptic currents during the recall process). We then examine our model in light of recent experimental findings (Michelmann, et al., 2016), where unique temporal signatures were detected for dynamic stimuli in the phase-angle time series. As such, we show the phase-angle time series of cortical regions at encoding (Figure 4Av, at 8Hz), where a phase-reset pattern coincides with the presentation of the pattern (red dotted box). Using representational-similarity-analysis (RSA), we can then compare this phase-reset pattern at encoding with the phase-angle time series at recall (Figure 4Ci), resulting in a similarity time-series that peaks at the time of the re-instatement of the encoded pattern in cortical regions (Figure 4Cii).

The experimental paradigm we chose to model investigated whether one could identify the reinstatement of a temporal stimulus from its unique temporal signature (Michelmann, et al., 2016). Therefore, we chose to present many patterns to our model with the aim of distinguishing each by its unique phasic signal. As such, all possible combinations of patterns consisting of 3 distinct events over a period of 1.5s were presented, with the first occurring at 0s to align stimulus onset and a further 2 events sampled at random from a set of uniformly spaced possibilities, where each of these 10 patterns were repeated 10 times (Figure 5, top panels; P1-P10). During encoding, we observed the familiar desynchronisation of the ongoing frequency (Figure 5Ai, 0-1.5s at the intrinsically generated 8Hz cortical rhythm) at stimulus onset, indicating that alpha desynchronisations potentially signify information processing (Klimesch, et al., 2007; Hanslmayr, et al., 2012).

**FIGURE 5.**
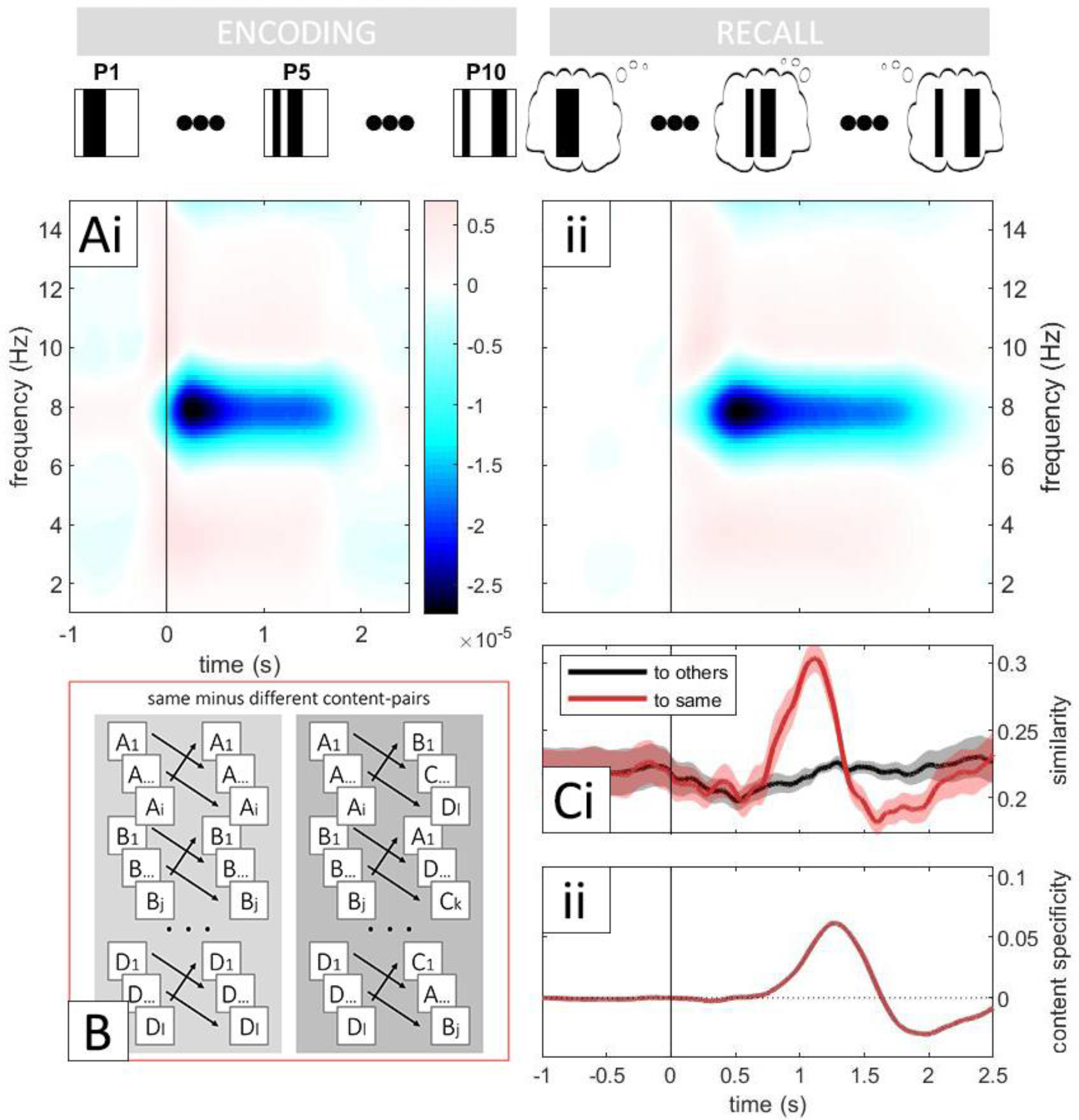
Content specificity over several patterns comprising 3 distinct events. Every possible combination of event timings is used over a period of 1.5s, where the 1^st^ occurs at 0s and the 2^nd^ & 3^rd^ are sampled at random from a set of uniformly spaced (.285 seconds) possibilities (top panels; P1-P10) and each being repeated 10 times. During encoding, a time-frequency analysis shows a desynchronisation of the ongoing oscillation across all patterns and trials (**Ai**; 0-1.5s at 8Hz, black vertical line indicates stimulus onset). As described in Figure 4, each pattern was replayed during recall, where a similar desynchronisation was also found (**Aii**). Representation-similarity-analysis (RSA) was then performed between encoding and recall phase-angle time series (**Ci**), where the average of similarity between the same patterns is indicated as a red line and similarities to other patterns as a black line (shaded regions indicate confidence intervals across trials and patterns, 50ms linear smoothing filter applied). As in the paradigm presented in Michelmann et al., 2016 (**B**; same minus different content pairs, adapted from Michelmann, et al., 2016), the difference between these similarities (**Cii**; same [**Ci**; red line] minus different [**Ci**; black line] content pairs, 500ms linear smoothing filter applied) was interpreted as evidence for content specific reinstatement of temporal patterns.

As shown in Figure 4, the recall phase entailed restarting the hierarchical synfire chain, such that any relevant bindings are reactivated in sequence. During this time, a pronounced alpha desynchronisation is also observable, building upon previous modelling work (Parish, et al., 2018) that indicates that this can predict both successful memory encoding and recall (Fell, et al., 2011; Hanslmayr & Staudigl, 2014; Hanslmayr, et al., 2012; Khader, et al., 2010; Klimesch, et al., 2005; Waldhauser, et al., 2016). A notable lag in cortical reinstatement (Figure 5Aii) indicates that the upwards direction of retrieval processes takes longer than the downwards direction of encoding processes, which is also indicated by experimental findings (Michelmann, et al., 2016; Griffiths, et al., 2019). Neurophysiologically, there are likely to be many neuronal layers for detecting and processing stimuli that would subsequently increase this recollection lag. In the model, however, this is mostly since binding units fire late in their respective temporal window (as indicated by Figure 4). This lag could be reduced within the model by increasing upwards directional weights such that binding units activated earlier in their window. However, this could have the undesirable effect of shifting activation forwards from encoding to recall, as events that are bound at a later point during the relatively broad window of our self-completing chain (∼200ms in length, ∼5Hz), might be replayed at an earlier point. In response to this, one could reduce the error by choosing a finer temporal dimension for the fastest, self-completing synfire chain, such that bindings would be encoded and recalled with more temporal precision. Such a notion might go some way to addressing why high-frequency gamma oscillations (>40Hz) are prevalent during episodic memory formation (Sederberg, et al., 2007). This line of reasoning has also been noted by Fell & Axmacher (2011) and other models of sequence encoding (Jensen, et al., 1996), who argue that gamma provides a fine-grain window of activation to maximise precise communication and learning.

Previous research indicates that a cortical alpha desynchronisation and subsequent phase-reset pattern might signify information flow (Haegens, et al., 2011; Hanslmayr, et al., 2012) and convey information content (Ng, et al., 2013; Schyns, et al., 2011), respectively. Here, we have shown some theoretical evidence for these findings, where a reduction in power (Figure 5Ai-ii) and phasic non-stationarity (Figure 4C) signifies the active representation of information. Further, experimental evidence has suggested that one can decode information content through analysis of the phase-angle time series, enabling the identification of dynamic stimuli through examination of their unique temporal signatures (Michelmann, et al., 2016). We next show that this position can also be supported theoretically through the use of computational simulations. By presenting many unique temporal patterns to our model (Figure 5; top panel), we can compare phasic signatures from encoding to retrieval between trials of the same pattern as well as to other patterns (Figure 5B; same minus different paradigm). Here, we use RSA, as in other research (Michelmann, et al., 2016), to compare these phase-reset patterns (as seen in Figure 4Cii), where a red line in Figure 5Ci indicates the average similarity between trials of the same patterns and a black line indicates the average similarity between trials of different patterns. The relatively flat line (<0.25) of the latter indicates that patterns were unique enough that on average, they did not resemble other patterns, whereas the peak in the former (>0.3, ∼1-1.5s) indicates that the phasic time-series of each pattern across trials was robust enough that on average, they resembled the same temporal signature. Taken together, the difference between these similarities (Figure 5Cii; same minus different) can indicate the content specific reinstatement of unique temporal patterns. The lag in the reinstatement of these patterns (∼1s) can partly be attributed to the lag discussed in the previous paragraph, and also resonates with the fact that the highest similarity occurs at the midpoint of comparable time-series (as seen in Figure 4Cii). The dip in performance after content specific recognition (∼1.8s), which is not present in the literature (Michelmann, et al., 2016), might be due to the nature of the patterns presented and to the relative stability of our intrinsically generated oscillations and circuitry in comparison to the brain. As such, it might be the case that there is some unusual systematicity due to our uniformly spaced patterns. This would mean that as the pattern is convolved, then the likelihood that the pattern exists in anti-phase to itself is higher, an unlikely occurrence in the brain due to highly distributed, dynamic and complex representations.

## DISCUSSION

We have here presented a novel neural network with three distinct mechanisms (see Figure 3 for model architecture). Together, these mechanisms enable us to bind discrete, observable events in time, though each mechanism allows us to explore a distinct set of hypotheses. We first instantiate a neo-cortical (NC) mechanism (Figure 3, top section), where intrinsic oscillations are generated at a resting alpha frequency through recurrent excitatory-inhibitory interactions. Here, we have implemented a winner-take-all mechanism in a “Mexican-hat” like topology, as has also been described in hierarchical models of vision, recognition and attention (Carpenter & Grossberg, 1987; Itti, et al., 1998; Reisenhuber & Poggio, 1999). By introducing an external event to our NC region, a subset of winning units that coded for that specific stimulus remained active whilst others were silent. During this time, the intrinsic frequency was de-synchronised (Figure 5Ai), consistent with the experimental hypothesis that oscillatory desynchronisations are due to increased neural activation (Haegens, et al., 2011; Hanslmayr, et al., 2011a), which occurs due to relevant representations becoming active at the expense of others (Klimesch, et al., 2007). In this way, we lend evidence to our primary hypothesis that desynchronised neural states indicate information flow within the brain (Hanslmayr, et al., 2012), as in previous modelling works (Parish, et al., 2018).

By adding a temporal dynamic to our model, we were also able to look at the phase-angle time series of the intrinsic frequency. In the literature, phasic patterns are thought to convey information content (Ng, et al., 2013; Schyns, et al., 2011), where one can even identify a stimulus by a unique temporal signature that can be used to later detect its re-instatement in memory (Michelmann, et al., 2016; Michelmann, et al., 2018; Michelmann, et al., 2019), a method that has also been applied in several recent MEG studies (Kornysheva, et al., 2019; Kurth-Nelson, et al., 2016; Lui, et al., 2019). The model lends theoretical evidence to both of these notions, showing that phase-reset patterns occur at a time when representations are active within our modelled NC region (Figure 4). Taking this further, we show how the model can store and recall a sequence of events, where representational-similarity-analysis (RSA) was used to show that phase-reset patterns at encoding and recall were sufficiently similar. Thus, we provide computational support for how phase can be important in conveying information content (Ng, et al., 2013; Schyns, et al., 2011). By presenting many sequences to our model, we were then able to show that one could on average detect a temporal stimulus by its phase-angle signature, so long as sequences were sufficiently unique (Figure 5Ci-ii), lending theoretical evidence to the experimental technique of using RSA to identify the content specific reinstatement of unique dynamic stimuli (as shown by Michelmann, et al., 2016).

Whilst we have successfully generated identifiable phase-reset patterns in our model, the best method of conveying information through this means remains unclear. Events were generated using a series of externally generated excitatory spike events, though we did not vary the number of spikes or the duration of sustained input between events. In further explorations of this parameter space, it would be interesting to demonstrate if the timing of these events relative to the up-down states of the intrinsic frequency influence whether they would be remembered or not, as has been found in the literature (Klimesch, et al., 2007). Within the model, this would hypothetically rely on the strength of cortical inhibitory projections, as well as the strength and duration of external events. Other models have also found that several regions, each with an independent intrinsic oscillation, can align in phase through the mediation of a coordinating pacemaker (Vicente, et al., 2008). As this is a proposed mechanism through which independent cortical columnal alpha oscillators are thought to align in phase, possibly through recurrent thalamo-cortical loops, it would be of further interest to instantiate a similar environment and assess whether phase can be robustly used to convey information content. One could then also explore whether a priming event might cause a general phase-reset in thalamic-cortical columns, as has been hypothesised to occur during the EEG P1 component of general recognition (Hanslmayr, et al., 2011b). This might ensure that cortical regions are pre-aligned in phase and thus optimally entrained to a given stimulus, such that any subsequent phase-reset patterns might be more consistent across trials and not as reliant on the intrinsic oscillatory conditions.

Episodic memories have an inherent temporal dynamic, whereby studies have suggested that our perception is not continuous but is rhythmically sampled in discrete alpha-frequency time-steps (Hanslmayr, et al., 2013; Landau & Fries, 2012; VanRullen, et al., 2007). It might therefore be the case that there is a more qualitative element to alpha desynchronisations. As neocortical and hippocampal gamma have both been found to be important when predicting successful memory encoding (Sederberg, et al., 2007), it might be the case that an alpha desynchronisation deregulates alpha-frequency inhibition in order for a gamma-frequency sequence to be allowed to activate. In this sense, our neocortex within the model might undergo learning to inherit a hippocampal sequence during encoding (or more likely, during sleep), as other models have shown is possible (Itskov, et al., 2011), even in the absence of a downstream feed-forward architecture. The deregulation of alpha inhibition might then enable the inherited cortical sequence to play in full, both decreasing alpha power and increasing respective gamma power.

The second mechanism within our model (Figure 3, middle section; binding pool) ensured that events were treated discretely such that they could be bound as independent components of a sequence. This method bears similarity to other models (Bowman & Wyble, 2007), where it was argued that the use of a binding-pool (BP) enables repetition of events to be represented. During development of the model, it was found that if bindings were direct from hierarchical time-keepers to cortical cells, then the cortical cells would activate upon every possible iteration of its bound hierarchical components. For example, if events were bound to the 3^rd^ part of the 1^st^ period (1,3) as well as the 1^st^ part of the 2^nd^ period (2,1), then during recall, the cortical assembly would activate at both the 1^st^ and 3^rd^ parts of both the 1^st^ and 2^nd^ period (1,1; 1,3; 2,1; 2,3; see “temporal conjunction errors” in Figure 2). The best prevention method of this is if there were only one layer of synfire chains, as has been hypothesised in other models (Itskov, et al., 2011), however, then the timed duration of the synfire sequence must be long enough to encode for the entire episodic sequence, which is often a criticism levelled against this type of model (Shankar & Howard, 2012). An investigation into these specific forms of temporal conjunction errors would give more insight into the brain’s adherence to the hierarchical encoding of time proposed here.

An interesting finding of our model is the effect of the “off switch” mechanism in the binding pool that operates to prevent spreading activation across the excitatory population (see Figure 3 for description). Here, it was found that a new event would not be encoded during this inhibitory pulse, as cortical impulses could not overcome the increased global inhibition. This is reminiscent of the hypothesised function of binding in other models (Bowman & Wyble, 2007), where it is thought that attentional deficits for events very close together in time arise as a consequence of the need to limit the temporal extent of the binding process (Wyble, et al., 2011). In our model, it is sufficient that the synaptic time constant from the “off-switch” cells to excitatory cells merely match that between excitatory cells, such that it quenches the possibility that activation ‘snowballs’ (see supplementary Figure S3 for more information on binding pool topology). However, if we are to relate this hypothesis (see Bowman & Wyble, 2007, & Swan & Wyble, 2014, for overview of binding errors and the attentional blink) to our model, then this time constant might be larger to act as a buffer between events, minimising binding errors as well as acting in its default role of preventing runaway activity. This might especially be necessary when encoding over very fine grain temporal dimensions, where a global inhibitory pulse could be useful in separating events close together in time, though it risks causing an attentional blink for those events that follow before inhibition subsides. Importantly, our model would also allow for multiple events to be encoded within the same binding location, possibly leading to order errors when replaying these events later, as has also been found in the literature for the encoding of events close together in time (Botella, et al., 1992; Wyble, et al., 2009). It would be of interest to simulate more of these paradigms, as in other models (Bowman & Wyble, 2007; Swan & Wyble, 2014), to determine whether these findings can be robustly linked to the functional limitations when modelling an iteration of a binding pool.

The third mechanism within our model (see Figure 1 & Figure 3A, bottom section) builds on the use of feed-forward synfire chains in the encoding of time. Due to the relatively short temporal constraints of our simulated episodic paradigm, we could have used a one dimensional synfire chain, as in other models (Goldman, 2009; Itskov, et al., 2011). However, we thought it appropriate to first address the main criticism levelled at such models (Shankar & Howard, 2012) that they are unlikely to be used for temporal encoding since the physical dimensions of the chain have to increase linearly with temporal requirements. This complexity issue is compounded when one considers that episodic sequences can be in the many 10s of seconds (Howard & Eichenbaum, 2013) and that there are likely many competing temporal lists (MacDonald, et al., 2013; Pastalkova, et al., 2008; Wood, et al., 2000). We address this issue by proposing a biophysical neural network model of persistent activation, which uses recursive feedforward and feedback connections to instantiate simultaneous temporal hierarchies. In this way, each hierarchical layer repeats, decreasing the computational complexity issue raised by Shankar & Howard, 2012, and events are encoded by their association to a point on multiple time-scales that together encode for a unique moment in time (see Figure 1 for full description of this clock-like analogy). In doing so, we also address another critisicm that has been levelled against recurrent mechanisms that enable persistent activity, raised by Goldman, 2009, where it has been argued that their use is unlikely to encode for long temporal durations due to the ever decreasing weights that would be required to sustain activity. This is overcome by an architecture of sequential groups of recurrent excitatory cells, each able to maintain persistent activation indefinitely in a ring-like architecture (see supplementary Figure S4). We also envisage a simple and compact neuronal assembly that can hypothetically be scaled up indefinitely for any number of chains of any length. This assembly ensures that recurrent activation can be maintained indefinitely with no adjustments to the parameter set, where a feedback signal is required in order to terminate activation and move the signal to the next neuronal group. This is with the exception of a necessarily self-completing (finest grain) dimension, which in the context of our model, must operate in the absence of feed-back. In reality, feedback for this dimension might originate from temporal or spatial boundaries due to large changes in environmental representation. Alternatively, the self-completing dimension could exist in a simpler architecture, like other models (Itskov, et al., 2011), where it might be contextualised by our hypothesised structure encoding for higher temporal dimensions.

It might also be the case that a signal could traverse chains at varying speeds dependent on the strength of the initial burst or levels of background noise, as other feed-forward synfire models have explored (Diesmann, et al., 1999; Kumar, et al., 2008). In this way, one could replay events at a faster rate, as has been observed during sleep (Diekelmann & Born, 2010). It might also be the case that some attentional mechanism can skip through scene-sized chunks, as indicated by recent findings (Michelmann, et al., 2019), where selective additional excitatory input focussed on a specific temporal dimension might compress memory replay until a point of interest is identified through an additional feedback mechanism.

We also consider our feedback update method between temporal hierarchies, where a parameter-dependent ‘handover’ period might exist (see Figure 2Aii). This raises several interesting questions, the first of which is how the now well-established phenomenon of order-errors (Bowman & Wyble, 2007; Wyble, et al., 2009) and conjunction errors (Botella, et al., 1992) might arise in episodic memory. Here, one can choose whether to maximise ‘what’ or ‘when’, minimising missed stimuli (object distinctiveness) or episodic binding errors (episodic distinctiveness), respectively. Secondly, a period of unresponsiveness during this ‘handover’ might provide a theoretical explanation for the functional significance of observable oscillatory dynamics, as gamma, theta and delta frequencies have all been found to be important for encoding and sequencing memories by modulating phases of learning. With respect to this idea, our model can be compared to a popular model of theta-gamma coupling (Jensen, et al., 1996), where items are segregated in time by gamma and theta time-steps. In our model, each respective synfire chain hierarchy might encode for a slower frequency, where items are bound in the respective up-states of gamma-theta coupling, and ‘handover’ periods at each hierarchy segregate memories into distinct episodes. This is suggestive of the oscillatory hierarchy that is thought to control neuronal excitability and stimulus processing (Lakatos, et al., 2005), as delta (1-4Hz) phase modulates theta (4-10Hz) amplitude, and theta phase modulates gamma (30-50Hz) amplitude. Further demonstrations of our hierarchical synfire chain model will focus on the idea that oscillations might have the functional role of segregating our temporal experiences. Alternatively, one could minimise these oscillatory blind-spots by instantiating a temporal population code, such that a continuous temporal episode can be fully covered by overlapping oscillators.

As our simulated paradigm is relatively short, we do not make full use of our timing mechanism. Future work could explore the full potential of this mechanism in encoding for very long periods of time, which could well provide an interesting neural substrate for processes such as the circadian rhythm (see Hofstra & de Weerd, 2008, for review). As this is a proof of concept mechanism, the structure of our temporal cell assembly also exists in a finely balanced parameter set. It would therefore be of interest to see if such a structure could reliably evolve through recurrent plastic connections, as has been shown to be the case for one dimensional synfire chains in another modelling study (Fiete, et al., 2010). Instead of persistent excitatory activation, these chains might similarly be organised through inhibitory heteroclinic orbits (Rabinovich, et al., 2006), or other such methods. Our main interest is in simulating some mechanism for the representation of a hierarchical encoding of time, and issues that might arise from such an approach.

## Supporting information

FIGURE 1 - synfire chains

FIGURE S4 - synfire chain topology

FIGURE S5 - encoding

FIGURE S6 - recall

## SUPPLEMENTARY MATERIALS

### 1. NEUROPHYSIOLOGY

Neurons in our model were simulated using Hodgkin-Huxley equations (as described in Equation 1). Here, membrane potential (*V_m_*) fluctuated over time according to intrinsic changes from specific ion channels for the leak current (I_L_; Equation 2), sodium (I_Na_; Equation 3) and potassium (I_K_; Equation 4), as well as extrinsic input from dendritic synapses (I_syn_) and applied direct current (I_DC_), where rate of change in the membrane potential (*d_Vm_/d_t_*) is modulated by the capacitance of the membrane (*c_m_*). Constants for the conductance of each channel dictate the rate of change of that channel 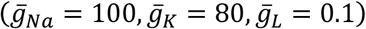, whilst the reversal potential drives change in each channel dependent on the membrane potential (*E*_*Na*_ = 50, *E_K_*> = −100, *E_L_* = −67). Voltage-dependent gates dictate the degree to which each channel is open, which are updated dependent on the current value of the voltage (*V_m_*) at time *t* (Equation 5). Changes in the gates *m*, *n* and *h* over time were modelled with opening rates of α_*x*_(*V_m_*) (Equations 6-8) and closing rates of β_*x*_(*V_m_*) (Equations 9-11).

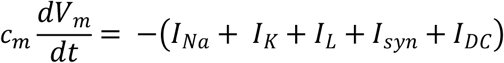

Equation 1 – The Hodgkin & Huxley model, 1952; channels for the sodium (Na), potassium (K), leak (L), dendritic synapses (syn) and direct current (DC) are modulated by the membrane capacitance (*cm*) to effect the membrane potential (*V_m_*).

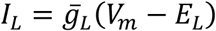

Equation 2 – The leak current; the driving force (difference between membrane potential (*V_m_*) and resting potential (*E_L_*)) multiplied by the conductance 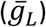.

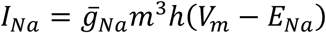

Equation 3 – The sodium (Na) current; the respective driving force multiplied by the respective conductance and the voltage-dependent gates *m* & *h*.

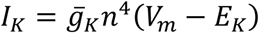

Equation 4 – The potassium (K) current; the respective driving force multiplied by the respective conductance and the voltage-dependent gate *n*.

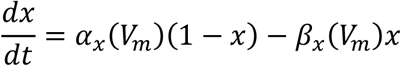

Equation 5 – Voltage-dependent (*V_m_*) gate update, where *x* denotes gate type.

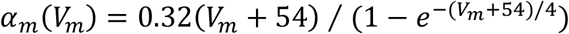

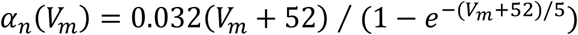

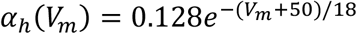

Equations 6, 7 & 8 – Voltage dependent (*V_m_*) opening of gates *m*, *n* & *h*, respectively.

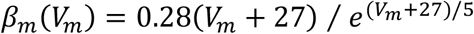

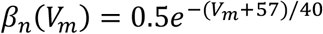

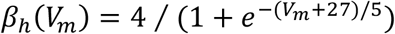

Equations 9, 10 & 11 – Voltage dependent (*V_m_*) closing of gates *m*, *n* & *h*, respectively. Spike events effect model neurons in the shape of an Alpha function to model post-synaptic-currents (Equation 12), which were multiplied by a weight (*w*). The synaptic time constant (τ*_s_*) dictates the amount of time it takes for the input to decay to 33% of its maximum value, where *t* indicates elapsed time since the spike event. All spike events have a delay of between 1-2ms to reach down-stream neurons.

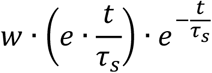

Equation 12 – Exponential Alpha function for a post-synaptic current (PSC).

Spike-time-dependent-plasticity (STDP) was enabled via an adapted calcium-based plasticity model (Equations 13-15; Graupner & Brunel, 2012). Synapses were represented by the variable *ρ*(t) (Equation 13), which existed in a state between 0-1 and was multiplied by specific weight variables. Synapses gravitate towards being active (ρ = 1) or inactive (ρ= 0) once they pass an attractor state (ρ_∗_). Synapses changed dependent on whether the amount of calcium *c* (t) in the neuron was over specific thresholds for long-term-potentiation (LTP; θ*_p_*) or long-term-depression (LTD; θ*_d_*), whereby LTP occurred at a rate γ*_p_* and LTD occurred at a rate γ*_d_* (Equation 13; *H* denotes the Heaviside function, which returns 0 or 1 if the expression within [] is below or above 0, respectively).

The amount of calcium at a synapse can be calculated by the summation of all spike events from pre (*i*) and post (*j*) synaptic neurons (Equation 14), where the Dirac delta function *δ*(*t* − *t_i_*) takes the elapsed time since each spike event to capture its decayed amount. In this model, an additional constraint was added to reduce multiple-spike calcium increases (Equation 15), such that the function *f*(*t_i_* − *t_i_*_−1_) takes the time between spike events and returns an exponentially fitted value that punishes short periods. Calcium is then summed subject to a multiplicative constant *C_pre_* for pre-synaptic spikes after a delay (*D* = 13*ms*), and a constant *c_post_* for post-synaptic spikes. Calcium also decayed exponentially over time (τ_*Ca*_ = 20*ms*).

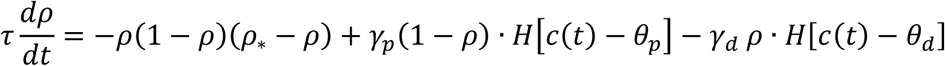

Equation 13 – Synaptic efficacy dynamics in calcium-based STDP (Graupner & Brunel, 2012).

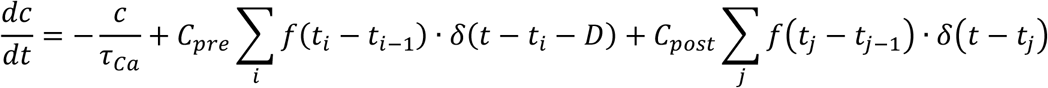

Equation 14 – Adapted calcium update equation.

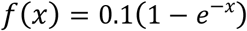

Equation 15 – Adaptive calcium increase equation.

### 2. TOPOLOGY & PARAMETERS (CHILDREN OF FIGURE 2)

**FIGURE S1.**
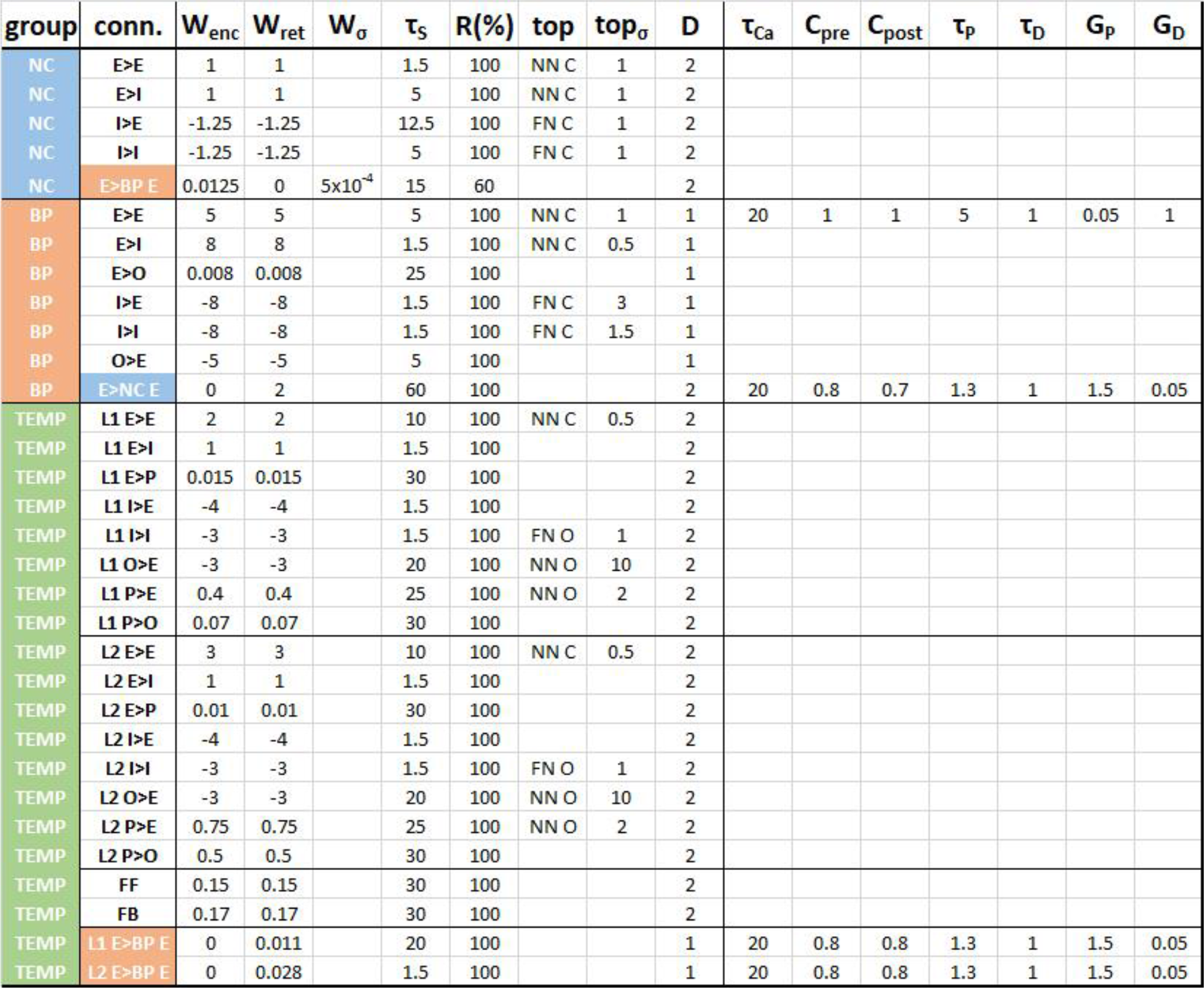
CONNECTIVITY & STDP PARAMETER SET. Parameter set of connectivity and spike-timing-dependent-plasticity (STDP) variables, where each column will be described in a left-right direction. The colour-coded far left column (group) designates originating region, whilst the next column to the right (conn.) details which connection is being represented. Here, directional connections (>) between excitatory (E), inhibitory (I), “off-switch” (O) and “propagation” (P) cells are described in each row. Temporal feed-forward (FF) and feed-back (FB) connections, as well as colour-coded intra-region connections, are also described. Simulations were comprised of two distinct phases, an encoding phase and a recall phase. Some intra-region weights were set to 0 to reflect the assumed presence of a neuro-modulatory effect that enables the switch between downwards encoding states (W_enc_) and upwards retrieval states (W_ret_). It is important to note that even if weights are here set to 0, the synapse variable *ρ* is still able to fluctuate between an active (1) and inactive (0) state if STDP is enabled on those connections. Zero mean Gaussian noise was also added to some weights (W_σ_). The synaptic time constant (τ_s_) determined the duration of dendritic post-synaptic-currents, and regions were connected with an R (%) chance to form a connection. Specific topologies (top.) were implemented between groups, nearest-neighbour (NN) connections between two banks of neurons were implemented as a Gaussian distribution centred over each neuron, whereas farthest-neighbour (FN) connections were implemented as 1 minus a Gaussian centred over each neuron, each with a specific standard deviation (top. σ). These topologies could be closed (C) in a circular loop, or open (O) in a sequential line. A fixed delay (D) from the spike of one neuron and the initiation of a post-synaptic-potential in the other was also implemented. As previously described, calcium dynamics were implemented where calcium decayed at a rate τ_Ca_, increased by constants C_pre_ & C_post_, where if thresholds for potentiation (τ_P_) or depression (τ_D_) were passed then the synapse would undergo potentiation or depression at rates of G_P_ or G_D_, respectively.

**FIGURE S2.**
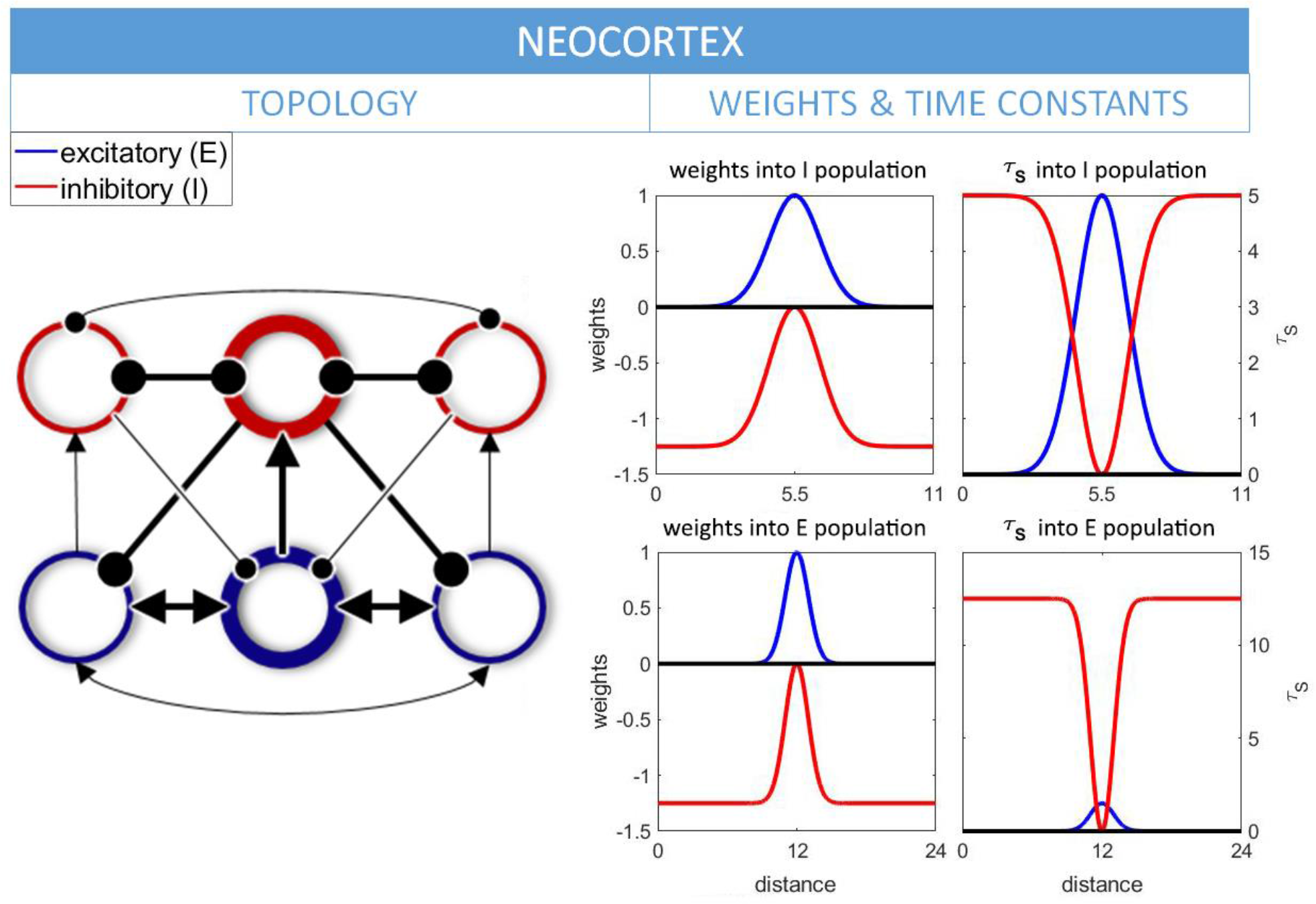
NEO-CORTEX TOPOLOGY. The neo-cortical region (NC) was modelled using a population of 35 neurons, 30% of which were inhibitory (I; red lines) and 70% excitatory (E; blue lines). These neurons were connected using surround inhibition to enable a winner-take-all dynamic (as shown in the left-hand panel), and as has commonly been used in hierarchical models of vision, recognition and attention (Carpenter & Grossberg, 1987; Itti, et al., 1998; Reisenhuber & Poggio, 1999). Excitatory neurons will oscillate at a steady state until some become more active, inhibiting their competitors in the process. To realise this, nearest- & farthest-neighbour connections were implemented between E and I banks of neurons, with specific weights (W) and synaptic time constants (τs) as shown in the right-hand panels. Here, colour-coded lines indicate input from E (blue lines) or I (red lines) sources, centred over the median neuron in the receiving group. In order for a winner-take-all topology to fall into an intrinsic, oscillatory steady-state, it was important to ensure activation could spread through the whole population before selective dynamics took hold. As such, the lower τ*_s_* for excitatory-excitatory connections (*τ_s_* into E population, blue line) allows excitatory activation to quickly spread through the population, before the slower-reacting, larger τ*_s_* for excitatory-inhibitory connections (*τ_s_* into I population, blue line) can in turn enable I neurons to clamp down on the E population. This causes oscillatory behaviour, where the frequency is set by the τ*_s_* for inhibitory-excitatory synapses (*τ_s_* into E population, red line; Brunel, 2000), thus here the frequency of idling activation is ∼8Hz, roughly approximating a resting alpha frequency. Maintaining a steady state oscillation requires that E neurons are constantly driven to activate by low level noise. To do this, each excitatory neuron is fed by a Poisson distribution of 1500 spike events per second (τ*_s_* = 1.5, *w* = 0.05), modelling activation from distant neurons and generating variation between neurons. The neo-cortical section of our model represents the flow of information, where the de-synchronisation and phase-angle time-series of alpha signals information processing (Hanslmayr, et al., 2012) and conveys information content (Canavier, 2015), respectively. Phase resets are thought to occur when there is a change in the scene during the presentation of dynamic stimuli (Michelmann, et al., 2016). In our model, scene changes are simulated as a strong pulse of Poisson distributed spike events (1000*Hz*, τ*_s_* = 1.5, *W* = 3), whose time course is multiplied by an Alpha function of τ*_s_* = 60*ms* (Equation 12). This input is fed into a random NC unit, where an overlapping Gaussian in the spatial dimension effects nearby units (*S. D.* = 1.5). Upon activation of this select group, respective inhibitory neurons also activate due to the nearest neighbour connections. These inhibit all other excitatory and inhibitory units, thus promoting the unimpeded activation of the selected group until the event driven stimulus dissipates. STDP was not implemented in our neocortex, as in a complementary learning systems framework (O’Reilly, et al., 2011), it was assumed that learning occurs very slowly in this region. As our model is a proof of principle ‘one-shot’ learning paradigm, slow STDP was not implemented here.

**FIGURE S3.**
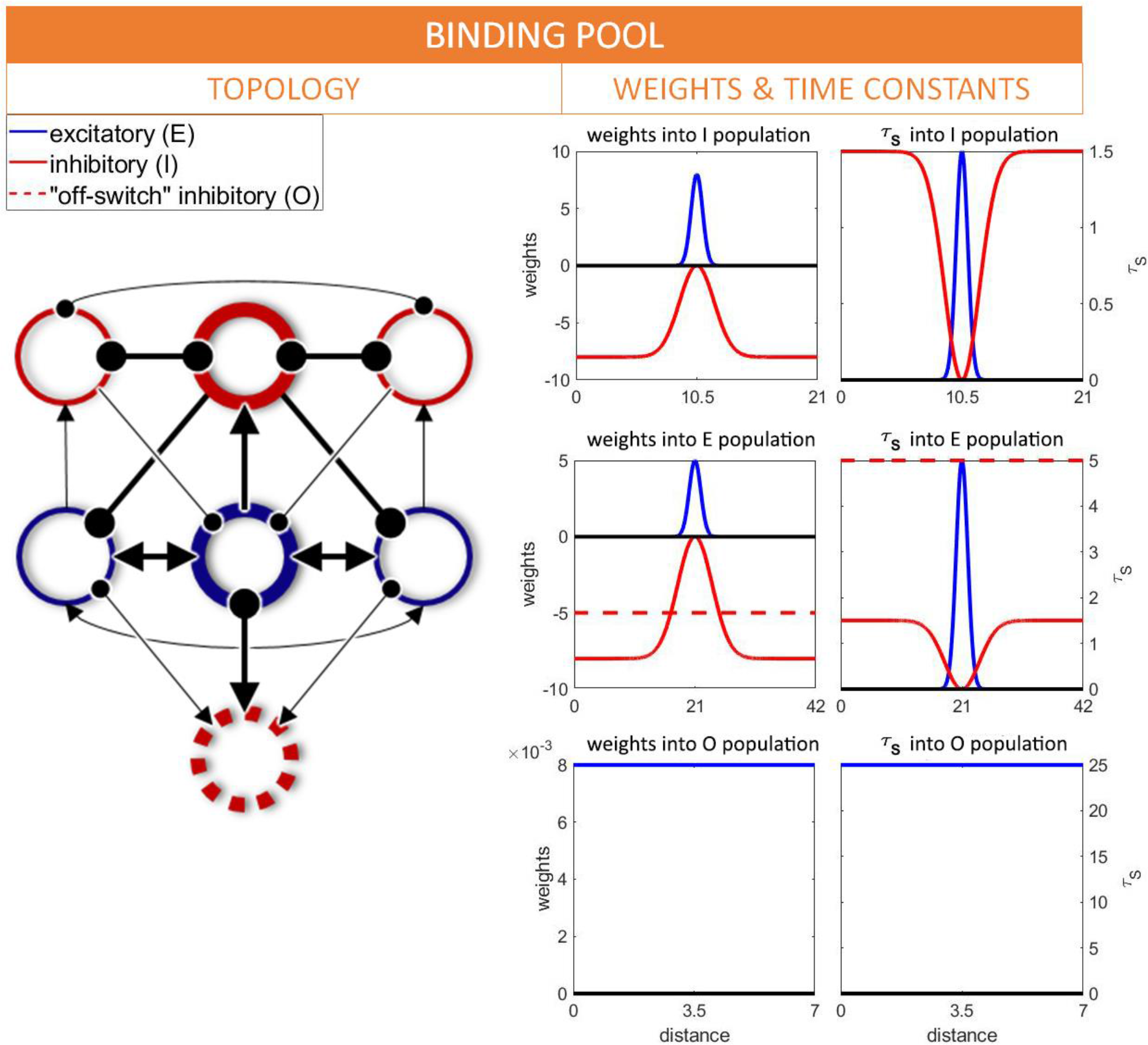
BINDING POOL TOPOLOGY. The binding pool (BP) was modelled using a population of 70 neurons, 30% of which are fast-inhibitory units (I; red lines), 10% “off-switch” inhibitory units (O; red dotted lines) and 60% excitatory (E; blue lines). Like the neo-cortex, surround inhibition was used here to promote a winner-take-all dynamic with nearest- & farthest-neighbour topologies. Here, the emphasis is placed on the low τ*_s_* for excitatory-inhibitory (*τ_s_* into I population, blue line) and inhibitory-excitatory (*τ_s_* into E population, red line) connections that allow competitive inhibition to spread quickly. A higher τ*_s_* for excitatory-excitatory connections (*τ_s_* into E population, blue line) ensures that initial E activity can survive the subsequent inhibitory response, thus giving preference to the earliest spikers due to the speed with which they can inhibit their competitors. Weights are much higher in the binding pool than in the neo-cortex, as this was found to be necessary to allow swift winner-take-all competition to take place. Binding pool units are much more selective, however, remaining silent until an event occurs, upon which one group will become active for a potentially indefinite period. To counter this, two additional mechanisms were implemented to promote selectivity and terminate activation. Firstly, STDP was added to synapses between binding pool units (see Figure S1 for STDP parameters), such that weights go down between active units. Here, calcium amplitudes are prevented from crossing the prohibitively high LTP threshold (θ*_p_*), yet cross the LTD threshold (θ*_d_*) during sustained activation. When this occurs, weights go down at a relatively fast rate (γ*_d_*). This weakens the active groups ability to participate in winner-take-all competition during successive events, creating a highly selective mechanism such that no one group of binding pool units will activate for any two events. Diminished internal weights also ensures that active neurons are less able to affect one another over time and activation within the group recedes. As this process occurs simultaneously with the long-term potentiation of other synapses within these active neurons, this mechanism effectively models a form of heterosynaptic LTD, as observed experimentally and in other models (Volgushev, et al., 2016). This process entails a balancing act where pre-existing synaptic pathways are diminished when new connections are strengthened through STDP. Alternatively, BP nodes could have been self-inhibited through sustained inhibition, as in other models that describe a binding-pool (Bowman & Wyble, 2007), or be modulated by an adaptive threshold mechanism, as in other models that instantiate a winner-take-all dynamic (Itskov, et al., 2011). Despite the currently employed mechanism, it is still possible for activation to spread through the network on occasions, disrupting the entire population. To this effect an “off-switch” (O unit; red dotted lines) was added, which is slow to activate but clamps down on the entire E population after a certain amount of activation has taken place. Parameters were chosen to ensure that input for the O units builds up slowly over a relatively long period, but once activated they send an immediate inhibitory pulse, where the τ*_s_* must be at least equal to that for excitatory-excitatory connections to mitigate any ongoing activation. For events to be transmitted to the binding pool, the neo-cortical excitatory bank connects to the binding pool excitatory bank (see Figure S1 for parameters). A large τ*_s_* and low weight discourage normal cortical oscillatory activity from triggering the binding pool, while encouraging the build-up of input from sustained spike-events that occur during cortical events. To maximise competition within the binding pool, randomness had to be implemented in NC-BP projections. To this effect, the banks connect via a random 15% of synapses, and weights vary according to a normal distribution. In order for events to be bound, STDP was implemented on the reverse BP-NC projections, where binding is encouraged with a fast LTP rate (γ*_p_*) and slow LTD rate (γ*_d_*). BP-NC connectivity was strong and long (τ*_s_* = 60, *W* = 2), to enable re-activation of NC events to occur on a similar time-scale as during encoding. Whilst this is an unrealistically large synaptic time-constant, this effectively models the accumulation of sustained activation through successive hippocampal-cortical pathways. One could also reasonably add a delay here due to the nature of this pathway, though this was not realised in the current model.

**FIGURE S4.** HIERARCHICAL SYNFIRE CHAIN TOPOLOGY (CHILD OF FIGURE 1) SEE VIDEO FILE CELL ASSEMBLY.MP4 As shown in Figure 1, hierarchical synfire chains (SC) are modelled as chains of sequentially connected groups existing within hierarchical layers, akin to a clock (Barnard, 2002; Friston, et al., 2018). Here, order denotes the position of the chain in the hierarchy, such that the slowest temporal dimension is termed the ‘highest-order’ chain and the fastest dimension the ‘lowest-order’ chain. Each group exists as a compact assembly of 8 units (see video for cellular assembly), 12.5% of which are inhibitory (I; red lines), 12.5% of which are “off-switch” inhibitory (O; red dotted lines), 12.5% are “propagation” excitatory (P; blue dotted lines) and 62.5% are excitatory (E; blue lines). We identify the need for two types of chains; higher-order chains that are terminated through feedback interactions with lower-order chains, and a lowest-order chain that must self-terminate due to there being no available sources of feed-back termination. In reality, we might expect feedback signals to originate from environmental changes, but in the framework of our model, we simply initiate a slightly different parameter range to encourage the propagation of activation in lowest-order chains (see Figure S1). Otherwise, each group in a higher-order chain provides feed-forward excitation to the first node in a lower order group (see * in above video), before receiving a terminating signal from the combined culmination of every existing lower-order chain (see ** in above video), upon which transient activity is propagated onto the next group. Within each group, E units exist in a ring like structure, each unit connecting in a loop to its nearest neighbour to promote persistent activation as described by Goldman, 2009. These all connect to the I unit in the group, which in turn connects to the E units and I units of every other group. This surround inhibition ensures only one group may be active at any one time, a key concept of time cells (Eichenbaum, 2014). P units enable propagation, receiving activation from their own group of E units and forwarding activation to the E units of groups either side (for ease of viewing, we show uni-directional flow in Figure S4, though in reality connections go bi-directional). This would in theory also enable reverse-playback of temporal sequences, though this is not simulated in the current work. All P units have feed-forward connections to the first E group in any existing lower-order chain (see Figure S1, FF connection), triggering the lower order chain to commence. P units also connect to the O unit within their group, where they either increase input to just below the spiking threshold for higher-order chains, or immediately activate O units in the lowest-order chain to enable self-termination (the key difference in the parameter space between temporal dimensions). All O units in higher-order chains then receive the summation of input from all of the P units in the final group of any existing lower order chain (see Figure S1, FB connection), which is only sufficient to push ready-to-fire O units to spike. It is important that the weight here is modified dependent on the number of layers in the hierarchy, such that *W* = *W*/*n*, where *n* denotes the number of feed-back connections from higher-order layers in relation to the current layer. Spiking O units then inhibit E units within their group, thus terminating the currently active group in the chain. This architecture enables P units to act as densely connected signal propagators, helping to maintain the input of groups of E units that it connects to, thus facilitating the signal to move onto the next E group in the chain, which it has been keeping in a ready-to-fire state. P units also perform feed-forward initiation (*) and feed-back termination (**), kick-starting the 1^st^ group of a lower-order chain via the former and terminating currently active groups in any higher-order chains via the latter. Within the context of this model, the highest-order temporal chain must be initiated by an external force, possibly attentional-related and fixed to the onset of a stimulus. This was implemented as an injection of a DC current to the first E unit in the first group lasting for 10ms (*W* = 0.15), which occurred 500ms before the onset of the first stimulus to allow the hierarchy to ramp up activation.

### 3. SIMULATIONS OF ENCODING AND RECALL (CHILDREN OF FIGURE 4)

**FIGURE S5.** ENCODING DISCRETE EVENTS IN TIME (CHILD OF FIGURE 4A) SEE VIDEO FILE ENCODING.MP4 The above video demonstrates the intra-cellular processes occurring during the encoding stage of our simulated paradigm (see Figure 4A). On the left-hand panels, we show normalised histograms of regional membrane voltages through time, with an overlapping Gaussian filter of width 20mV as a visual aide. This is done for each region, aligning with the raster plot of parent Figure 4; i.e. NC E (1^st^ row; blue histogram) & I (1^st^ row; red histogram) units, BP E (2^nd^ row; blue) & O (2^nd^ row; red) units, synfire chain (SC) layer 2 (L2; lowest-order chain) E (3^rd^ row; blue) & O (3^rd^ row; red) units, and SC layer 1 (L1; higher-order chain), E (4^th^ row; blue) & O (4^th^ row; red) units. Membrane voltage is shown on the x-axis, where spike events are visualised as a rapid movement in the positive direction and subsequent reset at the most negative end. In this way, one can visualise the internal dynamics of a group of neurons as they respond to inputs. SC regions (3^rd^ and 4^th^ rows) are further split into several overlapping histograms of varying shades, each corresponding to an individual group in the chain. The right-hand panels indicate intra-cellular calcium dynamics and weight changes, as defined by Equations 13-15 and the parameter sets defined in Figure S1. Here, calcium amplitudes (left-axis; red lines) increase in response to pre- & post-synaptic neuronal firing. If amplitudes are in between the thresholds for depression (red dashed line) and for potentiation (blue dashed line), then weights (ρ; right-axis, black lines) decrease, though if they surpass the potentiation threshold then weights (ρ) increase. Rows here indicate STDP enabled intra-regional connections (see Figure S1); i.e. BP->NC connections (1^st^ row), BP<->BP connections (2^nd^ row), SC L2 -> BP (3^rd^ row) and SC L1 -> BP (4^th^ row). Additionally, we show neocortical phase through time in the topmost panel. Describing from the bottom row upwards, one can see the effect of the initial DC burst being fed into a group of the SC L1 E units, as several rapid spike events are accompanied by a burst in calcium amplitudes from SC L1 to BP units (∼2000ms). One can then see how the active group in the SC L1 row inhibits its rival (indicated by shifts to the left of the voltage axis), whilst a regular sub-threshold calcium amplitude to BP units ramps up. There is also a very slight positive shift in one group of SC L1 O units, as it is being kept in a ready-to-fire state by its accompanying active E units. Next, at ∼2500ms, feed-forward initiation causes one group of SC L2 E cells to spike, along with the aforementioned effect of surround inhibition that maintains temporal context by inhibiting other SC groups (visualised as sharp movements to the left of the voltage axis). This is also accompanied by an increase in calcium amplitudes from SC L2 E units to BP E units to sub-threshold levels. One can see how calcium amplitudes are somewhat oscillatory, due to the ramping up of excitatory activation within successive groups. Due to the self-termination aspect of this layer of synfire chains, a single O unit can be seen to spike on occasion, causing the currently active group of SC L2 units to be inhibited, upon which the next group begins to spike. Eventually, the ready-to-fire O unit of SC L1 will also spike (∼4700ms) due to a feed-back termination signal from the final SC L2 P unit, thus causing activation in the successive SC L2 E group to ramp up over the subsequent 500ms. Now describing from the top row and downwards from the beginning of the video, one can see how NC units intrinsically oscillate. First, NC E units gradually shift in the positive direction until a cascade of spike-events occurs. This triggers their neighbouring I units to spike, which in turn inhibit the whole population of E units. Calcium amplitudes also periodically increase from BP E to NC E units during this rhythmic, post-synaptic spiking. Eventually, at ∼2500ms, an external event feeds spikes into a subset of NC E units. This causes a bi-modal distribution to occur in the membrane voltages of NC E & I units, as one group dominates the population (as seen in the parent raster plot of Figure 4A). The same phenomenon occurs in the highly selective BP E units, as they are triggered to fire by the additional NC input. When BP E units spike, calcium amplitudes in every row increase, as the binding pool is the glue that binds temporal position to content. This causes calcium to raise above the potentiation threshold in BP -> NC and both SC -> BP pathways, which in turn causes those directional weights (ρ; black line) to increase. This is with the exception of the BP <-> BP pathway, where calcium amplitudes are between potentiation and depression thresholds, causing a subset of weights to decrease from their previous maximal strength. Upon sufficient BP E activity, BP O units spike, triggering inhibition of the whole BP E layer and the termination of the encoding of that particular NC event.

**FIGURE S6.** RECALLING TEMPORAL SEQUENCES (CHILD OF FIGURE 4B) SEE VIDEO FILE RECALL.MP4 The above video demonstrates the intra-cellular processes occurring during the recall stage of our simulated paradigm (see Figure 4B). On the left-hand panels, we show normalised histograms of regional membrane voltages through time, with an overlapping Gaussian filter of width 20mV as a visual aide. As before, this is done for each region, aligning with the raster plot of the parent Figure 4. On the right-hand panels, we this time show synaptic inputs (I_syn_) that arrive into the respective E units of each region, whose resulting membrane voltages through time are shown in the left-hand panels. Here, blue shaded regions indicate excitatory input, whilst red shaded regions indicate inhibitory input. Describing from the bottom row upwards, one can once again see the effect of the initiating DC burst into SC L1 E units, though this time one can see that in the right-hand panel, ramping excitatory input maintains persistent activation in a feed-forward manner (blue shaded region, <2500ms), akin to Goldman, 2009, whilst inhibitory pulses perform surround inhibition to maintain temporal context (red shaded bursts, <2500ms), akin to Itskov, et al., 2011. At ∼2500ms, one can see resultant input from the activation of “propagation” (P) units, which have a longer *τ_s_* and therefore, a more sustained excitatory effect on E units (overlapping, darker blue shaded region). As these units initiate feed-forward activation of SC L2 E units, one can see how ramping input similarly builds up in this layer at this time. The self-termination nature of SC L2 E units becomes apparent at ∼2750ms, where a large inhibitory pulse terminates the currently active group. Importantly, the P unit here (signified by the overlapping, darker blue shaded region) enables transient activity to ramp up in the successive group due to the sustained input it provides, that overlaps both E groups. Once again, one can see the effects of this synaptic drive play out in the histogram of the membrane voltages. During this time, ramping synaptic input has been building up in BP E neurons due to persistent activation from SC L1 E units, due to the selectively strengthened synapses from the encoding phase. In addition to this, selective input also builds up from relevant SC L2 E units, though with a different time constant. The effect here is that the higher-order, slower synfire chain provides sustained input to keep bound units in a ready-to-fire state, whilst the lowest-order, faster synfire chain provides burst-like input that pushes bound units to fire, signalling the occurrence of an event in a hierarchical moment in time, much like recalling an event as occurring in the 10^th^ minute of the 2^nd^ hour. It is important that the activation of only one of these layers is not sufficient to trigger bound events, as this would cause temporal conjunction errors (as described in Figure 3). As before, the NC has been intrinsically oscillating at its designated frequency, due to low level background noise (shaded blue region, < 2500ms) that causes a periodic response to travel across the whole population. Here, large inhibitory synaptic inputs dictate how long it will take for E units to be able to recover and spike again, thus the inhibitory synaptic time-constant dictates the frequency of the intrinsically generated oscillation (Brunel, 2000). There is a reversal of directional weights between the BP <-> NC & SC <-> BP in the recall phase of the simulation (see parameters in Figure S2 for description of neuromodulators). This neuromodulation enables the directional flow of information and prevents recurrent activation between layers during encoding or recall. This is especially key to allow the binding pool to fulfil its dual purpose, as it is required to be highly sensitive to either cortical or temporal activation during encoding and recall, respectively. As such, this allows binding pool units to activate due to the concurrent activations of hierarchical temporal positions in the recall phase, which in turn triggers a large, sustained input in NC E units. As before, this causes a bimodal distribution of NC E membrane voltages, as selective units that were previously bound in time re-activate in the encoded temporal sequence. The effect of this can be seen in the phasic plot above.

## Notes

https://github.com/GP2789/The-Sync-fire-deSync-Model

## REFERENCES

1. Barnard, P., 2002. Mapping neural architecture to mental architecture and mental architecture to behavioural architecture. [Online] Available at: DOI: 10.13140/RG.2.2.36214.14401

2. Botella, J., Garcia, M. L. & Barriopedro, M., 1992. Intrusion patterns in rapid serial visual presentation tasks with two response dimensions. Perception & Psychophysics, 52(5), pp. 547–552.

3. Bowman, H. & Wyble, B., 2007. The Simultaneous Type, Serial Token Model of Temporal Attention and Working Memory. Psychological Review, 114(1), pp. 38–70.

4. Brown, G. D., Preece, T. & Hulme, C., 2000. Oscillator-Based Memory for Serial Order. Psychological Review, 107(1), pp. 127–181.

5. Brunel, N., 2000. Dynamics of Sparsely Connected Networks of Excitatory and Inhibitory Spiking Neurons. *Journal of Computational Neuroscience*, Volume 8, pp. 183–208.

6. Burgess, N. & Hitch, G., 1999. Memory for Serial Order: A Network Model of the Phonological Loop and its Timing. Psychological Review, 106(3), pp. 551–581.

7. Buzsáki, G., 2002. Theta Oscillations in the Hippocampus. *Neuron*, Volume 33, pp. 325–340.

8. Canavier, C. C., 2015. Phase-resetting as a tool of information transmission. *Current Opinion in Neurobiology*, Volume 31, pp. 206–213.

9. Carpenter, G. A. & Grossberg, S., 1987. A massively parallel architecture for a self-organizing neural pattern recognition machine. *Computer Vision*, Graphics, and Image Processing, 37(1), pp. 54–115.

10. Diekelmann, S. & Born, J., 2010. The memory function of sleep. Nature Reviews, 11(2), pp. 114–126.

11. Diesmann, M., Gewaltig, M.-O. & Aertsen, A., 1999. Stable propagation of synchronous spiking in cortical neural networks. *Nature*, Volume 402, pp. 529–533.

12. Eichenbaum, H., 2014. Time cells in the hippocampus: a new dimension for mapping memories. Nature Reviews, Volume 15, pp. 732–744.

13. Fell, J. & Axmacher, N., 2011. The role of phase synchronisation in memory processes. Nature Review Neuroscience, Volume 12, pp. 105–118.

14. Fell, J. et al., 2011. Medial Temporal Theta/Alpha Power Enhancement Precedes Successful Memory Encoding: Evidence Based on Intracranial EEG. Journal of Neuroscience, 31(14), pp. 5392–5397.

15. Fiete, I., Senn, W., Wang, C. & Hahnloser, R., 2010. Spike-time-dependent plasticity and heterosynaptic competition organize networks to produce long scale-free sequences of neural activity. Neuron, 65(4), pp. 563–576.

16. Fries, P., 2005. A mechanism for cognitive dynamics, neuronal communication through neuronal coherence. Trends in Cognitive Science, Volume 9, pp. 474–480.

17. Friston, K. et al., 2018. Deep Temporal Models and Active Inference. Neuroscience and Biobehavioural Reviews, Volume 90, pp. 486–501.

18. Goldman, M. S., 2009. Memory without Feedback in a Neural Network. *Neuron*, Volume 61, pp. 621–634.

19. Graupner, M. & Brunel, N., 2012. Calcium-based plasticity model explains sensitivity of synaptic changes to spike pattern, rate, and dendritic location. PNAS, 109(10), pp. 3991–3996.

20. Griffiths, B. J. et al., 2019. Directional coupling of slow and fast hippocampal gamma with neocortical alpha/beta oscillations in human episodic memory. PNAS, 116(43), pp. 21834–21842.

21. Haegens, S. et al., 2011. α-Oscillations in the monkey sensorimotor network influence discrimination performance by rhythmical inhibition of neuronal spiking. PNAS, 108(48), pp. 19377–19382.

22. Hanslmayr, S., Gross, J., Wolfgang, K. & Shapiro, K., 2011b. The role of alpha oscillations in temporal attention. Brain Research Reviews, pp. 331–343.

23. Hanslmayr, S. & Staudigl, T., 2014. How brain oscillations form memories - a processing based perspective on oscillatory subsequent memory effects. *Neuroimage*, Volume 85, pp. 648–655.

24. Hanslmayr, S., Staudigl, T. & Fellner, M.-C., 2012. Oscillatory power decreases and long-term memory: the information via desynchronization hypothesis. Frontiers in Human Neuroscience.

25. Hanslmayr, S. et al., 2013. Prestimulus Oscillatory Phase at 7 Hz Gates Cortical Information Flow and Visual Perception. *Curr Biol. Elsevier Ltd*, Volume 23, pp. 2273–2278.

26. Hanslmayr, S., Volberg, G., Wimber, M. & Raabe, K.-H. T., 2011a. The relationship between brain oscillations and BOLD signal during memory formation: a combined EEG-fMRI study. *Journal of Neuroscience*, Volume 31, pp. 15674–15680.

27. Hodgkin, A. L. & Huxley, A. F., 1952. A QUANTITATIVE DESCRIPTION OF MEMBRANE CURRENT AND ITS APPLICATION TO CONDUCTION AND EXCITATION IN NERVE. *Journal of Physiology*, Issue 117, pp. 500–544.

28. Hofstra, W. A. & de Weerd, A. W., 2008. How to assess circadian rhythm in humans: A review of literature. Epilepsy & Behvaior, Volume 13, pp. 438–444.

29. Howard, M., Viskontas, W., Shankar, K. H. & Fried, I., 2012. Ensembles of human MTL neurons ‘‘jump back in time’’ in response to a repeated stimulus. *Hippocampus*, Volume 22, pp. 1833–1847.

30. Howard, W. M. & Eichenbaum, H., 2013. The Hippocampus, Time, and Memory Across Scales. Journal of Experimental Psychology, 142(4), pp. 1211–1230.

31. Itskov, V., Curto, C., Pastalkova, E. & Buzsaki, G., 2011. Cell Assembly Sequences Arising from Spike Threshold Adaptation Keep Track of Time in the Hippocampus. Journal of Neuroscience, 31(8), pp. 2828–2834.

32. Itti, L., Koch, C. & Niebur, E., 1998. A model of saliency-based visual attention for rapid scene analysis. IEEE Transactions on Pattern Analysis and Machine Intelligence, 20(11), pp. 1254–1259.

33. Jensen, O., Idiart, M. & Lisman, J., 1996. Physiologically Realistic Formation of Autoassociative Memory in Networks with Theta/Gamma Oscillations: Role of Fast NMDA Channels. Learning & Memory, pp. 243–256.

34. Jensen, O. & Mazaheri, A., 2010. Shaping functional architecture by oscillatory alpha activity: gating by inhibition. Frontiers in human neuroscience, 4(186).

35. Khader, P., Jost, K., Ranganath, C. & Rosler, F., 2010. Theta and Alpha oscillations during working-memory maintenance predict successful long-term memory encoding. Neuroscience Letters, 468(3), pp. 339–343.

36. Klimesch, W., Barbel, S. & Sauseng, P., 2005. The Functional Significance of Theta and Upper Alpha Oscillations. Experimental Psychology, 52(2), pp. 99–108.

37. Klimesch, W., Sauseng, P. & Hanslmayr, S., 2007. EEG alpha oscillations: The inhibition–timing hypothesis. Brain Research Reviews, 53(1), pp. 63–68.

38. Kornysheva, K. et al., 2019. Neural Competitive Queuing of Ordinal Structure Underlies Skilled Sequential Action. Neuron, 101(6), pp. 1166–1180.

39. Kropff, E. & Treves, A., 2008. The Emergence of Grid Cells: Intelligent Design or Just Adaptation?. *Hippocampus*, Volume 18, pp. 1256–1269.

40. Kumar, A., Rotter, S. & Aertsen, A., 2008. Conditions for Propagating Synchronous Spiking and Asynchronous Firing Rates in a Cortical Network Model. The Journal of Neuroscience, 28(20), pp. 5268–5280.

41. Kurth-Nelson, Z., Economides, M., Dolan, R. & Dayan, P., 2016. Fast Sequences of Non-spatial State Representations in Humans. Neuron, 91(1), pp. 194–204.

42. Lakatos, P. et al., 2005. An Oscillatory Hierarchy Controlling Neuronal Excitability and Stimulus Processing in the Auditory Cortex. J Neurophysiology, Volume 94, pp. 1904–1911.

43. Landau, A. & Fries, P., 2012. Attention Samples Stimuli Rhythmically. *Curr Biol. Elsevier Ltd*, Volume 22, pp. 1000–1004.

44. Lui, Y., Dolan, R., Kurth-Nelson, Z. & Behrens, T., 2019. Human Replay Spontaneously Reorganizes Experience. Cell, 178(3), pp. 640–652.

45. MacDonald, C. J., Carrow, S., Place, R. & Eichenbaum, H., 2013. Distinct hippocampal time cell sequences represent odor memories in immobilized rats. *Journal of Neuroscience*, Volume 33, pp. 14607–14616.

46. Mauk, M. D. & Buonomano, D. V., 2004. The Neural Basis of Temporal Processing. Annual Reviews Neuroscience, Volume 27, pp. 307–340.

47. McNaughton, B. L. et al., 2006. Path integration and the neural basis of the ‘cognitive map’. *Nature Reviews Neuroscience*, Volume 7, pp. 663–678.

48. Michelmann, S., Bowman, H. & Hanslmayr, S., 2016. The Temporal Signature of Memories: Identification of a General Mechanism for Dynamic Memory Replay in Humans. PLoS Biology, 14(8).

49. Michelmann, S., Bowman, H. & Hanslmayr, S., 2018. Replay of Stimulus-specific Temporal Patterns during Associative Memory Formation. Journal of cognitive neuroscience, 30(11), pp. 1577–1589.

50. Michelmann, S., Staresina, B., Bowman, H. & Hanslmayr, S., 2019. Speed of time-compressed forward replay flexibly changes in human episodic memory. Nature Human Behaviour, 3(2), p. 143.

51. Moser, E., Kropff, E. & Moser, M.-B., 2008. Place Cells, Grid Cells, and the Brain’s Spatial Representation System. Annual Reviews Neuroscience, Volume 31, pp. 68–89.

52. Ng, B., Logothetis, N. & Kayser, C., 2013. EEG Phase Patterns Reflect the Selectivity of Neural Firing. *Cereb Cortex*, Volume 23, pp. 389–398.

53. O’Reilly, R., Bhattacharyya, R., Howard, M. & Ketz, N., 2011. Complementary Learning Systems. Cognitive Science, pp. 1–20.

54. Parish, G., Hanslmayr, S. & Bowman, H., 2018. The Sync/deSync Model: How a Synchronized Hippocampus and a Desynchronized Neocortex Code Memories. Journal of Neuroscience, 38(14), pp. 3428–3440.

55. Pastalkova, E., Itskov, V., Amarasingham, A. & Buzsaki, G., 2008. Internally generated cell assembly sequences in the rat hippocampus. *Science*, Volume 321, pp. 1322–1327.

56. Rabinovich, M. I., Huerta, R. & Varona, P., 2006. Heteroclinic Synchronization: Ultrasubharmonic Locking. *Physical Review Letters*, Volume 96.

57. Reisenhuber, M. & Poggio, T., 1999. Hierarchical models of object recognition in cortex. *Nature Neuroscience*, Volume 2, pp. 1019–1025.

58. Rolls, T. E. & Mills, P., 2019. The Generation of Time in the Hippocampal Memory System. Cell Reports, Volume 28, pp. 1649–1658.

59. Schreiner, T. et al., 2018. Theta Phase-Coordinated Memory Reactivation Reoccurs in a Slow-Oscillatory Rhythm during NREM Sleep. Cell Reports, 25(2), pp. 296–301.

60. Schyns, P., Thut, G. & Gross, J., 2011. Cracking the Code of Oscillatory Activity. *PLoS Biology*, Volume 9.

61. Sederberg, P. B. et al., 2007. Hippocampal and neocortical gamma oscillations predict memory formation in humans. *Cerebral Cortex*, Volume 17, pp. 1190–1196.

62. Shankar, K. & Howard, M., 2012. A Scale-Invariant Internal Representation of Time. *Neural Computation*, Volume 24, pp. 134–193.

63. Staresina, B. P. et al., 2016. Hippocampal pattern completion is linked to gamma power increases and alpha power decreases during recollection. *eLife*, Volume 5, p. e17397.

64. Staudigl, T. & Hanslmayr, S., 2019. Reactivation of neural patterns during memory reinstatement supports encoding specificity. Cognitive Neuroscience, 10(4), pp. 175–185.

65. Staudigl, T., Vollmar, C., Noachtar, S. & Hanslmayr, S., 2015. Temporal-Pattern Similarity Analysis Reveals the Beneficial and Detrimental Effects of Context Reinstatement on Human Memory. *Journal of Neuroscience*, Volume 35, pp. 5373–5384.

66. Swan, G. & Wyble, B., 2014. The binding pool: A model of shared neural resources for distinct items in visual working memory. Attention Perception & Psychophysics, Volume 76, pp. 2136–2157.

67. Tsao, A. et al., 2018. Integrating time from experience in the lateral entorhinal cortex. *Nature*, Volume 561, pp. 57–62.

68. VanRullen, R., Carlson, T. & Cavanagh, P., 2007. The blinking spotlight of attention. Proc Natl Acad Sci, Volume 104, pp. 19204–19209.

69. Vicente, R. et al., 2008. Dynamical relaying can yield zero time lag neuronal synchrony despite long conduction delays. Proc. Natl Acad. Sci. USA, Volume 105, pp. 17157–17162.

70. Volgushev, M. et al., 2016. Partial Breakdown of Input Specificity of STDP at Individual Synapses Promotes New Learning. Journal of Neuroscience, 36(34), pp. 8842–8855.

71. Waldhauser, G. T., Braun, V. & Hanslmayr, S., 2016. Episodic memory retrieval functionally relies on very rapid reactivation of sensory information. Journal of Neuroscience, 36(1), pp. 251–260.

72. Wimber, M. et al., 2012. Rapid Memory Reactivation Revealed by Oscillatory Entrainment. Current Biology, 16(21), pp. 1482–1486.

73. Wood, E. R., Dudchenko, P., Robitsek, R. J. & Eichenbaum, H., 2000. Hippocampal neurons encode information about different types of memory episodes occurring in the same location. *Neuron*, Volume 27, pp. 623–633.

74. Wyble, B., Bowman, H. & Nieuwenstein, M., 2009. The attentional blink provides episodic distinctiveness; sparing at a cost. J Exp Psychol Hum Percept Perform, 35(3), pp. 787–807.

75. Wyble, B., Potter, M. C., Bowman, H. & Nieuwenstein, M., 2011. Attentional episodes in visual perception. J Exp Psychol Gen, 140(3), pp. 488–505.

76. Yaffe, R. et al., 2014. Reinstatement of distributed cortical oscillations occurs with precise spatiotemporal dynamics during successful memory retrieval. PNAS, 111(52), pp. 18727–18732.

